# SMARCA2 is an essential and potent cofactor for a specific subset of the glucocorticoid response in A549 cells

**DOI:** 10.1101/2025.11.13.688306

**Authors:** Schuyler M Melore, D Dewran Koçak, Keith Siklenkla, Graham D Johnson, Courtney Williams, Luke C Bartelt, Angela G Jones, Alejandro Barrera, Revathy Venukuttan, Charles A Gersbach, Timothy E Reddy

**Affiliations:** University Program in Genetics & Genomics, Duke University, Durham, NC, USA; Department of Biostatistics & Bioinformatics, Duke University, Durham, NC, USA; Department of Biomedical Engineering, Duke University, Durham, NC, USA; Altius Institute for Biomedical Sciences, Seattle, WA, USA

## Abstract

Glucocorticoids are a widely used, potent class of anti-inflammatory drugs that modulate the expression of hundreds of genes across the genome. Although the glucocorticoid response is primarily carried out by the glucocorticoid receptor (NR3C1, a.k.a. GR), there are many glucocorticoid receptor co-factors that are also essential to the downstream effects. To identify novel factors necessary for the glucocorticoid gene expression response, we used a genome-wide CRISPR screen in A549 lung adenocarcinoma cells. In that screen, we knocked out every gene in the human genome, and measured the effect of expression of the glucocorticoid-induced leucine zipper (GILZ), a classic glucocorticoid-response gene. We identified two chromatin remodeling proteins, SMARCA2 and BPTF, that are essential for GILZ expression. We then evaluated the genome-wide effects of SMARCA2 and BPTF on glucocorticoid-mediated gene expression. BPTF had a highly specific role in the glucocorticoid response, affecting the expression of only a handful of genes, and having virtually no effect on dexamethasone-induced changes in chromatin accessibility. However, SMARCA2 was necessary for 27% of dexamethasone-induced transcriptional changes (152 genes), and ∼7% of dexamethasone-induced changes in chromatin accessibility (586 regions of the genome). Genomic regions with SMARCA2-dependent changes in chromatin accessibility were characterized by high dexamethasone-induced regulatory activity in a massively parallel reporter assay, and dexamethasone-induced increases in transcription factor binding and chromatin states. Taken together, these data suggest that SMARCA2 is critical for chromatin remodeling at a specific set of genomic regions with high regulatory activity, which in turn drive changes in expression for many glucocorticoid-responsive genes.

## Introduction

Glucocorticoids are among the most widely prescribed small molecule pharmaceuticals due to their strong anti-inflammatory activity^1^. Their long-term use is severely limited, however, by a wide range of adverse effects that impact many bodily systems^2^. A longstanding pharmaceutical goal is to develop selective glucocorticoid agonists that elicit the strong anti-inflammatory effects of glucocorticoids, but with fewer side effects. Those efforts have been successful in modulating the overall potency of glucocorticoid activity; but have not been successful in selectively eliciting subsets of the overall response^1,3^.

At a molecular level, glucocorticoid responses occur when small molecules bind glucocorticoid receptors sequestered in the cytosol. Upon binding, the receptor is transported into the nucleus where it acts as a transcription factor^4^. While the glucocorticoid receptor is centrally important in that pathway, the downstream effects on gene expression involve interactions with many other proteins including other transcription factors (e.g. AP-1, CEBPB), chromatin remodeling factors (e.g. SMARCA4), nuclear receptor co-factors (e.g. NCOA and NCOR proteins)^4–8^. Those interactions could be alternative pharmaceutical targets that could be used to more selectively control the glucocorticoid response.

Here, we sought to identify novel protein factors contributing to the glucocorticoid response and the subset of gene expression responses that depend on those factors. To do so, we completed a high-throughput genome editing screen to identify proteins that are necessary for the glucocorticoid receptor to activate expression of the glucocorticoid-induced leucine zipper (GILZ, a.k.a. *TSC22D3*) gene. GILZ was one of the first glucocorticoid receptor targets identified, and it plays a key role in the anti-inflammatory effects of the GR^9,10^.

We identified and validated many known and novel genes that are necessary for the glucocorticoid receptor to upregulate GILZ. We especially focused on the chromatin remodeling factor SMARCA2 both because its role in the glucocorticoid-response is not well-characterized, and because it presents a clear mechanistic hypothesis. We show that a specific subset of the glucocorticoid receptor-dependent gene expression and chromatin remodeling require SMARCA2. In particular, glucocorticoid-induced changes in the expression of genes regulating apoptosis were enriched among SMARCA2-dependent gene expression changes. Together, these results demonstrate both a novel role for SMARCA2 in controlling the glucocorticoid receptor response; and more generally that focusing on genetic manipulation of co-factors is an effective strategy for selectively targeting subsets of the transcriptional response to glucocorticoids.

## Results

### A genome-wide knockout screen reveals necessary factors for glucocorticoid-induced GILZ expression

In order to conduct a screen for regulators of the glucocorticoid response, we adopted the A549 cell line as a model system, whose response to glucocorticoids we have characterized extensively^11^. Using this dataset, we evaluated genes via RNA-seq data for those with high overall expression levels as well as high fold changes in response to dexamethasone. GR binding was also evaluated by ChIP-seq to confirm that the genes are regulated in cis. This procedure led to the nomination of three genes (*ANGPTL4, TFCP2L1*, and *TSC22D3*), which were then evaluated using antibody staining. *TSC22D3* showed the strongest GC-induced staining and thus was chosen as the reporter for the genetic screen. *TSC22D3* is also one of the first discovered and most widely-studied glucocorticoid responsive genes. The protein product plays a key role in mediating the anti-inflammatory effects of glucocorticoids^9,10^.

To perform the screen, we used a pooled CRISPR/Cas9 knockout screening library targeting 19,114 genes in a pool of A549 cells^12,13^. A549 cells are a human lung carcinoma epithelial cell line that is commonly used for studying glucocorticoid responses^14–18^. Cells were transduced with the Brunello lentiviral CRISPR library, and after eight days were given a four hour treatmentment of 100 nM dexamethasone (dex), a synthetic glucocorticoid, or equal volume ethanol as a vehicle control. Cells were then harvested, stained using an anti-GILZ antibody, and sorted into high and low expression bins using FACS **(Fig. 1A)**. We then used MAGeCK to identify target genes whose sgRNAs were enriched in cells with the highest and lowest 10% GILZ expression for both the Dex- and vehicle-treated samples.

**Figure 1.**
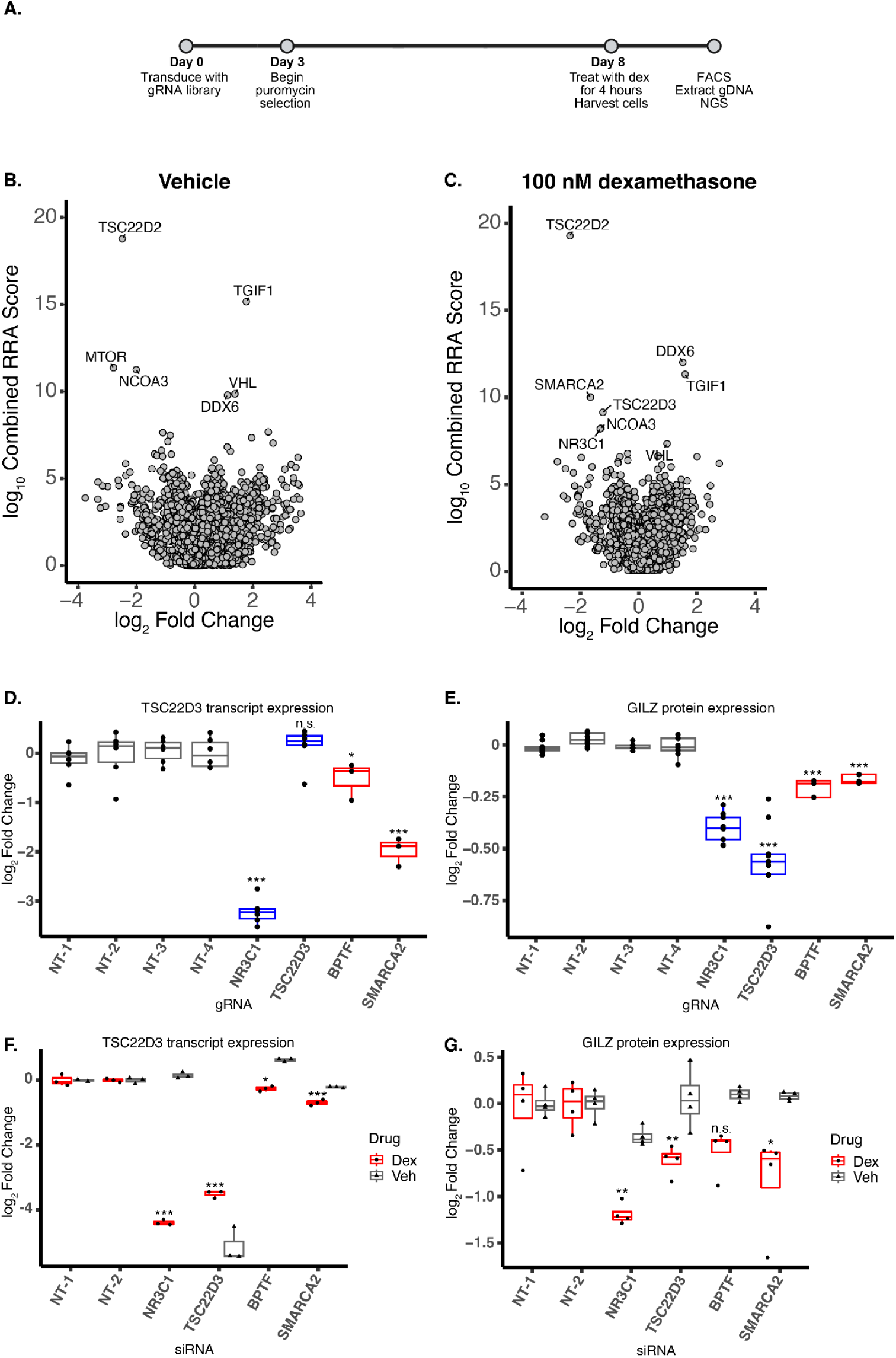
A genome-wide CRISPR screen to discover necessary factors for glucocorticoid-induced GILZ Expression. **A.** CRISPR screening timeline. Cells were transduced with the Brunello genome-wide CRISPR screening library at low MOI, selected with puromycin, and passaged for eight days. They were then treated with 100 nM dexamethasone or 0.1% ethanol as a vehicle control for four hours, stained with an anti-GILZ antibody, and sorted into high and low expression bins containing the top and bottom 10% GILZ-expressing cells.. gDNA was collected from cells, gRNA sequences were amplified, and enrichment in high and low bins vs. the bulk population was determined (n = 5 replicates per drug condition). **B,C.** Differential gRNA enrichment in the high and low GILZ expression bins in **B.** the vehicle control condition and **C.** the 100 nM dexamethasone treatment condition. Log_2_ fold changes are an average of high bin vs. bulk unsorted and low bin vs. bulk unsorted. Negative sign indicates a gRNA was depleted from the high bin and enriched in the low bin. (n = 5 replicates per drug condition) **D,E.** Validation of top gRNAs from the screen using **D.** qPCR or **E.** flow cytometry to determine the impact of knocking out the target gene on glucocorticoid-induced GILZ expression (n = 3 independent transductions). * p < 0.05, ** p < 0.01, *** p <0.001, one-tailed student’s t-test. Samples transduced with non-targeting control gRNAs are shown in gray, positive control gRNAs targeting *NR3C1* and *TSC22D3* are in blue, and gRNAs targeting *SMARCA2* and *BPTF* in red. **F,G.** Validation of top hits from the screen using siRNAs to knock down the genes of interest and assess their impact on glucocorticoid-induced GILZ expression using **F.** qPCR (n = 2 to n = 3 independent transfections) or **G.** flow cytometry (n = 4 independent transfections) * p < 0.05, ** p < 0.01, *** p <0.001, one-tailed student’s t-test

We identified three genes whose knockout impacted GILZ expression levels in dex-treated cells but not in cells treated with vehicle control: *NR3C1* (the glucocorticoid receptor), *TSC22D3* (the gene encoding GILZ), and *SMARCA2* **(Fig. 1B-C)**. SMARCA2 is a component of the mammalian SWI/SNF chromatin remodeling complex, which is composed of >10 subunits, including one of two mutually exclusive ATPases, SMARCA2 (also known as BRM) or SMARCA4 (also known as BRG1), that make up the catalytic subunit^19^. This complex has been previously shown to play a critical role in mediating the glucocorticoid response, and the GR has been shown to directly interact with several subunits^17,18,20–32^. However, the vast majority of those studies have focused on the role of SMARCA4-, not SMARCA2-containing complexes.

The specific roles of SMARCA2 and SMARCA4 in the cellular response to glucocorticoids appears to be highly cell type and context dependent. For example, in breast cancer and murine mammary adenocarcinoma cell models, SMARCA4 is necessary for a subset of the the glucocorticoid response by facilitating chromatin remodeling and GR binding at certain regions of the genome^22,23,28–32^. Another study used engineered HEK293T cells to identify proteins that interact with GR and the closely related androgen receptor (AR). That study found both GR and AR interact with SMARCA4, but only found evidence for AR interacting with SMARCA2 in that model^20^. In contrast, studies in a uterine cancer cell line and in an adrenal carcinoma cell line showed that SMARCA2 and GR together strongly activated a GR-responsive reporter after dex treatment, while GR alone could not activate the same reporter^25,27,33^. Lastly, studies in A549 cells, which harbor a 23 nt homozygous deletion in SMARCA4 rendering it non-functional, have shown that SMARCA2 interacts with GR and is necessary for dex-induced changes in transcription and chromatin accessibility at a handful of well-characterized dex-responsive loci, including *TSC22D3*^17,18,26,34^. Based on those results, we hypothesize that SMARCA2 can mediate glucocorticoid-associated chromatin remodeling genome-wide in cells that do not express SMARCA4.

We independently validated the effects on dex-induced *TSC22D3* transcription levels and GILZ protein levels for our top hits as well as several gRNAs with varying effects using qPCR and FACS **(Fig. 1D-E and Supp. Fig. 1A-D)**. We additionally validated these effects for *SMARCA2*, *BPTF*, *TSC22D3*, and *NR3C1* using siRNA knockdown. BPTF (bromodomain PHD finger transcription factor) is a key component of the NURF chromatin remodeling complex, and has the ability to direct the complex to different genomic loci through binding to histone marks. BPTF was not initially identified as a mediator of dex-induced GILZ expression in the screen since only one of the four gRNAs targeting it showed an effect. However, that gRNA showed an effect on both dex-induced *TSC22D3* expression levels and GILZ protein levels in the initial validation, leading us to prioritize BPTF for further followup. These experiments confirmed the effects we observed in the screen, where *NR3C1* and *TSC22D3* knockout and knockdown result in a complete loss of dex-induced GILZ expression while *SMARCA2* and *BPTF* knockout and knockdown results in a partial loss of dex-induced GILZ expression **(Fig. 1F-G, Supp. Fig. 1E-G)**. Of note, cells harboring the *TSC22D3* gRNA had similar *TSC22D3* mRNA levels to cells harboring a non-targeting gRNA, but much lower GILZ protein levels. We interpret that as the cells producing a gene-edited *TSC22D3* mRNA that does not produce a functional protein.

### SMARCA2-dependent transcriptional responses to dex treatment

We first sought to test the hypothesis that SMARCA2 and BPTF contribute to a specific subset of the genomic response to glucocorticoids. To do so, we measured changes in gene expression genome-wide in A549 cells after knockdown of *SMARCA2* or *BPTF* expression. As a positive control, we knocked down expression of the glucocorticoid receptor itself (*NR3C1*); and as a negative control, we delivered siRNAs designed to not target any human genes (i.e. non-targeting). After two days of knockdown, we treated cells for two hours with either 100 nM dexamethasone or 0.1% ethanol as a vehicle control. We then measured genome-wide changes in gene expression using RNA-seq.

Based on those RNA-seq results, knockdown was highly efficient for all targeted genes. *NR3C1* expression was reduced by > 90%; *SMARCA2* expression was reduced by 83%; and *BPTF* expression was reduced by 70% **(Supp. Fig 2)**. In the non-targeting control condition, the glucocorticoid response was similar to the responses reported previously. Specifically, we identified 1,245 dex-responsive genes with a non-targeting siRNA at a false discovery rate of 5%. Those responsive genes include canonical dex-induced genes such as *ERRFI1*, *ANGPTL4*, and *TSC22D3* as well as dex-repressed genes such as *IL11*, *IER2*, and *IER3* **(Fig. 2A, Supp. Fig. 3A-B)**^14,17,35–37^. We also found roughly equal proportions of dex-activated genes (54%) and dex-repressed genes (46%), as reported previously^11^.

**Figure 2.**
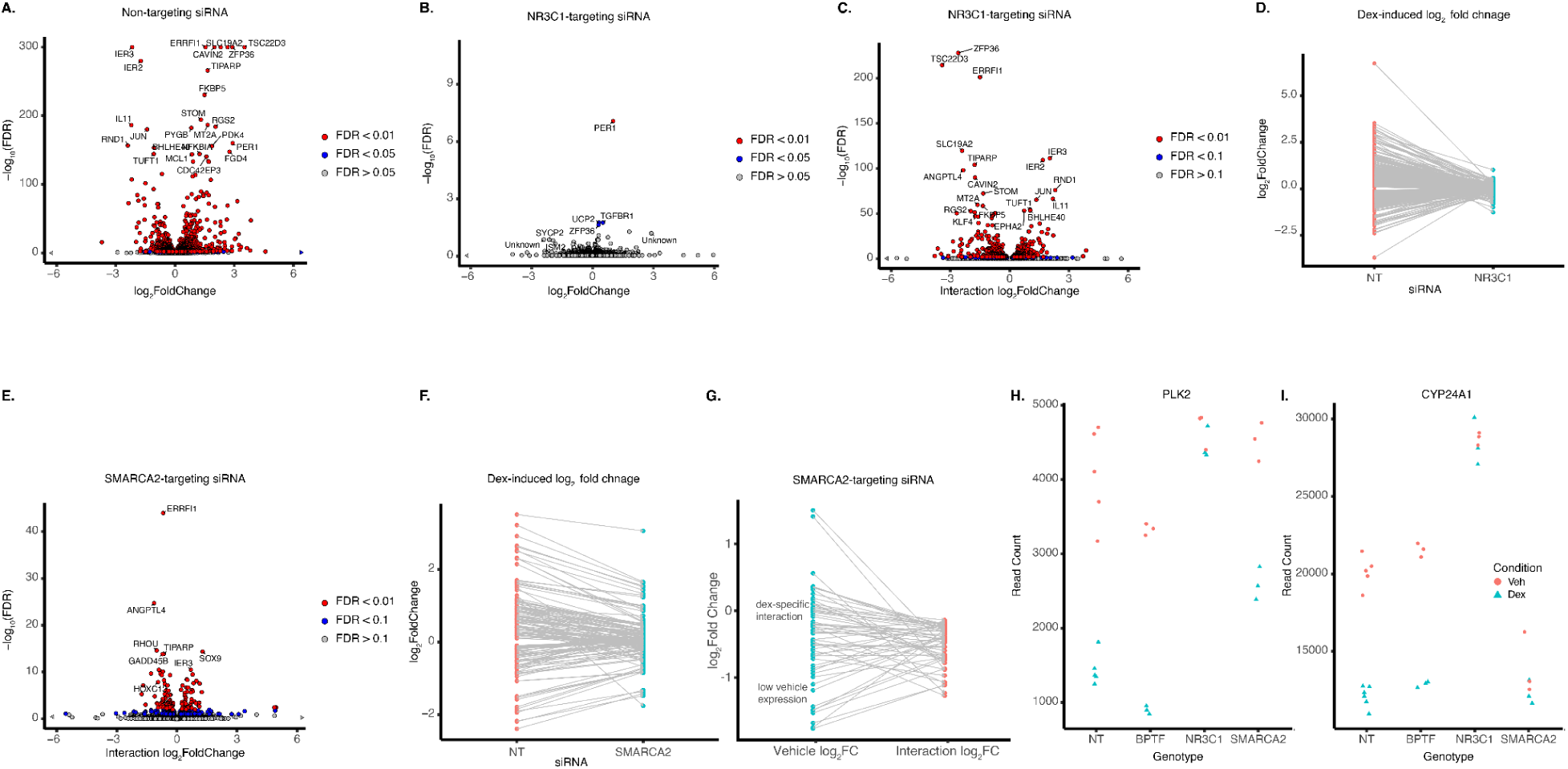
SMARCA2-Dependent Transcriptional Responses to Dexamethasone. **A.** Dexamethasone-induced transcriptional changes in cells transfected with a non-targeting siRNA. Cells were treated with either 100 nM dex or an ethanol vehicle control for two hours after which RNA was harvested for sequencing (n = 5 to n= 6 independent replicates per drug treatment condition). Fold change for dex vs. control treatment is shown. **B.** Dexamethasone-induced transcriptional changes in cells transfected with an *NR3C1*-targeting siRNA. Cells were treated with either 100 nM dex or an ethanol vehicle control for two hours after which RNA was harvested for sequencing (n = 3 independent replicates per drug treatment condition). **C.** Interaction effects for cells transfected with an *NR3C1-*targeting siRNA. The fold changes in this plot represent the difference between the fold change observed in NR3C1 knockdown cells and the fold change observed in non-targeting control cells. **D.** Fold changes after dex treatment for genes with GR-dependent transcription changes in cells transfected with non-targeting siRNA (left) or *NR3C1*-targeting siRNA (right) **E.** Interaction effects for cells transfected with *SMARCA2*-targeting siRNA. The fold changes in this plot represent the difference between the fold change observed in *SMARCA2* knockdown cells and the fold change observed in non-targeting control cells. **F.** Fold changes after dex treatment for genes with SMARCA2-dependent transcription changes in cells transfected with non-targeting siRNA (left) or *SMARCA2*-targeting siRNA (right) **G.** For the n = 59 SMARCA2-dependent, dex-repressed genes, this plot shows the log_2_ fold change compared to non-targeting control cells in the vehicle control condition (left) or the interaction log_2_ fold change (right). **H-I.** Normalized counts for SMARCA2-dependent, dex-repressed genes in dex and vehicle treatment conditions (n = 3 replicates per drug condition). **H.** For *PLK2*, *SMARCA2* knockdown does not affect baseline expression and dampens the repressive effect of dex treatment. **I.** For *CYP24A1*, *SMARCA2* knockdown decreases baseline gene expression, and dex treatment has no subsequent effect.

Knockdown of the glucocorticoid receptor itself nearly entirely ablated the glucocorticoid response. In a previous study, we detected a diminished glucocorticoid response in A549 cells with 50% knockdown of *NR3C1* expression^14^. We therefore hypothesized that the >90% knockdown achieved in this study should greatly reduce the glucocorticoid response. After knockdown of the glucocorticoid receptor, we detected only four dex-responsive genes that we also found in the non-targeting control response. Further, the response of those four genes was diminished compared to the non-targeting siRNA condition. **(Fig. 2B, Supp. Fig. 3A,C).**

*SMARCA2* and *BPTF* knockdown had similarly small effects on the overall glucocorticoid response. Overall, genes responded in the same direction with and without *SMARCA2* or *BPTF* knockdown, and overall the responses were strongly correlated (R > 0.8 for *SMARCA2* and *BPTF* knockdown compared to non-targeting control) **(Supp. Fig. 3D-E, Supp. Fig. 4)**. In addition, the median effect of dexamethasone treatment was similar between non-targeting and *SMARCA2* or *BPTF* knockdown. In the non-targeting condition, glucocorticoid-induced and -repressed genes had a median effect size of 1.37-fold and 0.78-fold respectively. After *SMARCA2* and *BPTF* knockdown, those responses were moderately attenuated. The median effect of induced-genes was reduced to 1.21-fold and 1.27-fold, respectively; and the median response of repressed genes was reduced to 0.86-fold and 0.84-fold respectively **(Supp. Fig. 5)**. Of the genes that are dex-responsive in the non-targeting condition, 502 and 640 were significantly dex-responsive after SMARCA2 or BPTF knockdown, respectively **(Supp. Fig. 3A, D-E)**. The reduced number of significantly dex-responsive genes and the modestly attenuated effect sizes after *SMARCA2* and *BPTF* knockdown raises the possibility that SMARCA2 and BPTF are required for a subset of the glucocorticoid response; however other explanations such as differences in statistical power to detect effects may also explain the differences.

**Figure 3.**
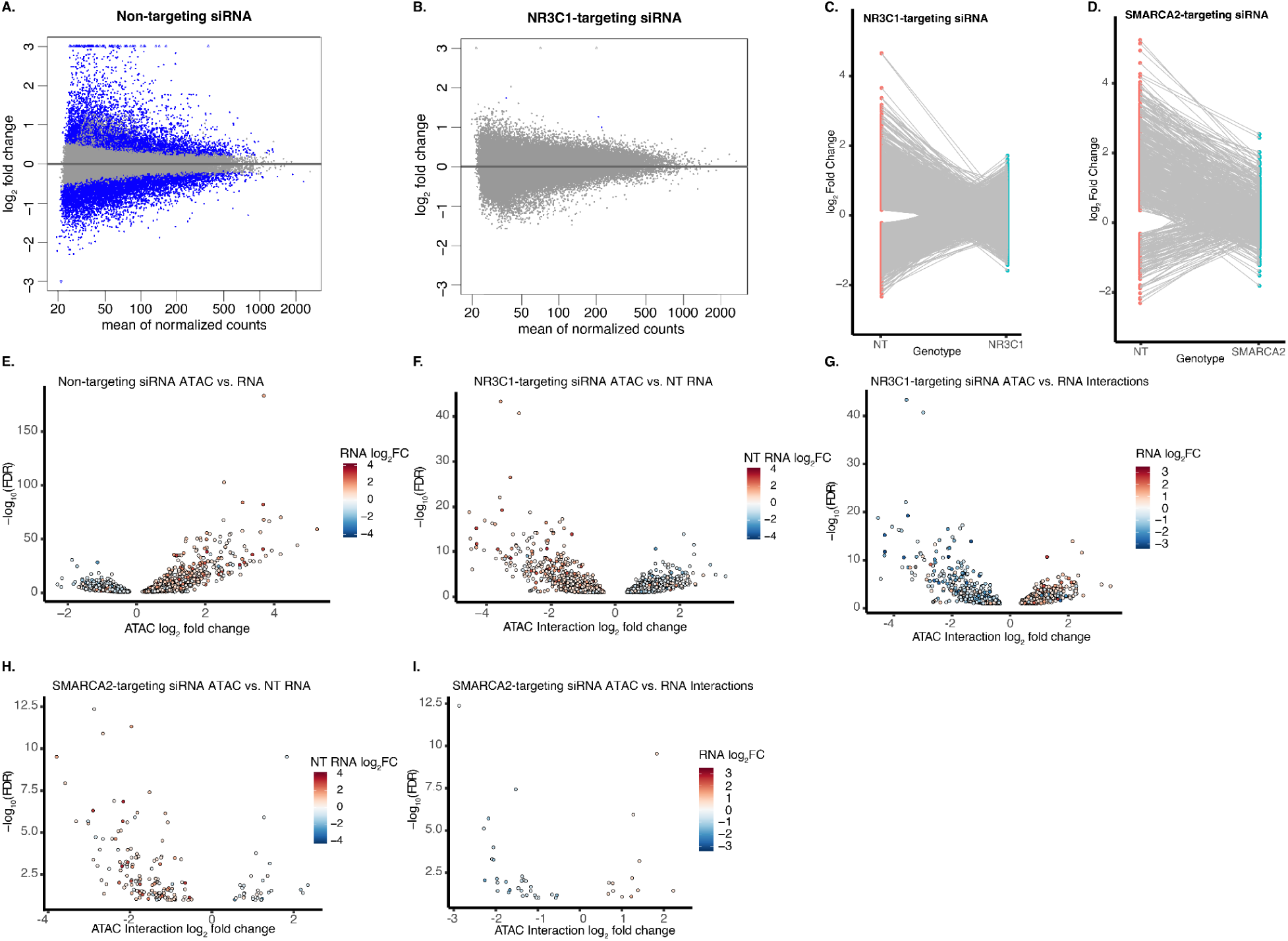
SMARCA2-dependent chromatin accessibility changes after dex treatment. **A.** MA plot showing dex-dependent changes in chromatin accessibility for cells transfected with a non-targeting siRNA (n = 6 independent transfections per drug condition). Cells were transfected with a non-targeting siRNA, and 48 hours later were treated with either 100 nM dexamethasone or a vehicle control for two hours before cells were harvested and ATAC-seq was performed. **B.** MA plot showing dex-dependent changes in chromatin accessibility for cells transfected with an *NR3C1*-targeting siRNA (n = 3 independent transfections per drug condition) **C,D.** Line plots showing the dex-induced log_2_ fold change for regions of the genome that had a significant interaction term for **C.** *NR3C1* or **D.** *SMARCA2*. Dex-induced log_2_ fold change in cells transfected with non-targeting siRNA is on the left, and dex-induced log_2_ fold change for those same regions, but in cells with the target gene knocked down, is shown on the right. **E.** Comparison of dex-dependent transcriptional responses and chromatin accessibility changes in cells transfected with a non-targeting siRNA. Volcano plot shows ATAC-seq log_2_ fold change for each genomic region that exhibited significant dex-induced chromatin accessibility changes and whose nearest gene exhibited significant dex-induced transcriptional response. The fill of each point indicates RNA-seq log_2_ fold change for the closest gene to that genomic region. **F-I.** Comparison of dex-dependent transcriptional responses and chromatin accessibility changes for *NR3C1* knockdown cells **F**. Volcano plot shows regions of the genome that exhibit GR-dependent changes in chromatin accessibility after dex treatment (significant interaction term for ATAC-seq) whose nearest gene was dex-responsive. The fill of each point indicates RNA-seq log_2_ fold change in the cells transfected with the non-targeting siRNA for the closest gene to that genomic region **G.** Volcano plot shows regions of the genome that exhibit GR-dependent changes in chromatin accessibility after dex treatment whose nearest gene exhibited an GR-dependent, dex-induced transcriptional change. The fill of each point indicates RNA-seq interaction log_2_ fold change for the closest gene to that genomic region. **H**. Volcano plot shows regions of the genome that exhibit SMARCA2-dependent changes in chromatin accessibility after dex treatment (significant interaction term for ATAC-seq) whose nearest gene was dex-responsive.The fill of each point indicates RNA-seq log_2_ fold change in the cells transfected with the non-targeting siRNA for the closest gene to that genomic region. **I.** Volcano plot shows regions of the genome that exhibit SMARCA2-dependent changes in chromatin accessibility after dex treatment whose nearest gene exhibited an SMARCA2-dependent, dex-induced transcriptional change. The fill of each point indicates RNA-seq interaction log_2_ fold change for the closest gene to that genomic region

Therefore, to rigorously test whether SMARCA2 or BPTF are required for the dex-responsiveness of specific genes, we fit a negative binomial linear regression model for each gene. In each model, we modeled the effect of knockdown as an interaction term between knockdown and dex treatment. We then identified genes for which there was both a significant dex-responsive effect in the non-targeting condition at a 5% FDR; and a significant interaction with the knockdown at a 10% FDR. After knockdown of the glucocorticoid receptor itself, 572 dex-responsive genes had significant interaction terms; and the magnitude and direction of those interactions were consistent with the receptor’s essential role in the response **(Fig. 2C-D, Supp. Fig 6A-B)**.

For *SMARCA2* knockdown, we identified 178 significant interaction terms. About 90% of those interactions (N = 152) reduce the dex response compared to cells treated with the non-targeting siRNA **(Fig. 2E-F, Supp. Fig 6A,C)**. Based on these results, we define a set of 152 SMARCA2-dependent glucocorticoid responses in A549 cells, representing 27% of dex-responsive genes in those cells. SMARCA2 knockdown substantially reduced dex-responsiveness across those genes (**Fig. 2F**). Our results replicated previously identified SMARCA2-dependent responses such as the dex-activated genes *ANGPTL4*, *MT2A*, and *CDKN1C*, as well as the dex-repressed genes *AMIGO2* and *BHLHE40*^17,26^. We also find that SMARCA2 knockdown impacts more dex-activated (61%) genes than dex-repressed genes (39%), which echoes previous work that has shown that SMARCA4 knockdown in murine mammary adenocarcinoma cells impacts more dex-activated than dex-repressed genes^30^. Though the effect of *SMARCA2* knockdown was overall smaller than the effect of *NR3C1* knockdown, there was no significant difference between the percent of the glucocorticoid response explained by *SMARCA2* or *NR3C1* interaction terms (p = 0.43, paired t-test). Additionally, *SMARCA2* knockdown did not have a significantly different effect on dex-activated vs dex-repressed SMARCA2-dependent genes (p = 0.82, paired t-test). **(Fig. 2F, Supp. Fig. 5, Supp. Fig. 6C).** For BPTF, there were 25 significant interaction effects, of which 21 reduced the magnitude of the glucocorticoid response. The overall magnitude of BPTF interactions terms were much smaller than for SMARCA2 knockdown **(Supp. Fig. 6D)**. Together, those results indicated that both SMARCA2 and BPTF contribute to the glucocorticoid response of specific genes, and that the effects of SMARCA2 are more numerous and greater than that of BPTF. Notably, the second-most significant BPTF interaction was for *TSC22D3*, confirming the results of our screen and further supporting the possibility the BPTF has very specific effects on glucocorticoid responses.

To investigate if SMARCA2 might control specific biological functions that the GR elicits, we tested for overrepresented gene functions using Panther^38,39^. As expected, multiple gene ontology categories related to the glucocorticoid response were significantly enriched, such as cellular response to corticosteroid stimulus and response to stress (FDR = 7.61×10^-4^ and 1.81×10^-7^, respectively). We also found that 44 (29%) of the 152 SMARCA2-dependent responses were for genes involved in apoptosis or the regulation of apoptosis. For example, the gene ontology categories for regulation of programmed cell death and for regulation of apoptotic processes were both significantly enriched (FDR = 9.37×10^-12^ and 4.27×10^-11^, respectively). As specific example genes, the pro-apoptotic factors *FOXO3* and *THBS1* were less induced in *SMARCA2* knockout cells, while negative regulators such as *IER3* and *NUAK2* were less repressed. Together, these results suggest that SMARCA2 may specifically contribute to apoptotic effects of glucocorticoids.

One possible explanation for the effects of SMARCA2 on glucocorticoid responses is that changes in gene expression with *SMARCA2* knockdown either reduce the potential range of expression for those genes, or make glucocorticoid responses more difficult to detect. Across all conditions, *SMARCA2* knockdown explains 38% of the variation in global gene expression according to a principal components analysis **(Supp. Fig. 6E)**. SMARCA2-dependent, dex-induced genes had similar expression in the SMARCA2-knockdown compared to cells treated with the non-targeting control siRNA in the vehicle control condition [mean log_2_(fold change) = 0.04, p = 0.59]. However, SMARCA2-dependent, dex-repressed genes had lower expression in *SMARCA2* knockdown compared to cells treated with the non-targeting siRNA in the vehicle control condition [mean log_2_(fold-change) = -0.37, p = 0.000]). In contrast, we found that the expression of these same dex-repressed genes in the *NR3C1* knockdown was more similar to cells treated with the non-targeting control siRNA [mean log_2_(fold-change) = -0.12, p = 0.03].

We hypothesized that some effects occur through a glucocorticoid-independent mechanism where steady-state gene expression is downregulated in response to *SMARCA2* knockdown and that renders the gene unresponsive to dex treatment. In such cases, we expect that *SMARCA2* knockdown would reduce expression in vehicle-control treatment, and that reduction in expression would explain most of any observed interaction effect. Overall, ∼40% of dex-repressed genes match that description, with repression in the vehicle control condition explaining the loss of dex-responsiveness with *SMARCA2* knockdown **(Figs. 2G-I)**.

### SMARCA2 is required for increasing chromatin accessibility at a specific subset of dex-responsive genomic regions

SMARCA2 is a key member of the SWI/SNF complex responsible for remodeling chromatin at gene regulatory elements. Previous work has shown that, while most GR binding sites are in chromatin that is already accessible before glucocorticoid exposure, those sites have increased chromatin accessibility after GR binding. The same studies also showed that SMARCA4-containing SWI/SNF complexes are necessary for chromatin remodeling at a subset of GR binding sites^22,30–32^. One potential mechanism for SMARCA2-dependent glucocorticoid responses is that loss of SMARCA2 prevents or limits chromatin remodeling at GR binding sites, and in doing so limits binding of GR and co-factors. To test that hypothesis, we measured changes in chromatin accessibility using ATAC-seq. As before, we did so in A549 cells after transfecting with either non-targeting (NT) siRNAs or siRNAs targeting *SMARCA2*, *NR3C1*, or *BPTF*; and then subsequently treating with either 100 nM dex or a vehicle control for 2 hours before cells were harvested for ATAC-seq.

Overall, there were 13,437 genomic regions that had significant changes in chromatin accessibility (FDR < 10%) after dex treatment **(Fig. 3A, Supp. Fig. 7A)**. *NR3C1* knockdown almost entirely ablated those chromatin responses, with only four regions of the genome identified as significantly dex-responsive **(Fig. 3B, Supp. Fig. 7A)**. In both the *SMARCA2* and *BPTF* knockdown conditions, a substantial fraction of the dex responsive regions in the non-targeting conditions still had a significant change in chromatin accessibility (N = 813 and N = 3,051), indicating the neither knockdown fully ablated the glucocorticoid response **(Supp. Fig. 7A-C)**. That reduced number of chromatin responsive sites in the knockdown conditions could be due to SMARCA2 or BPTF playing an essential role in the glucocorticoid response, but could also be due to differences in statistical power to detect changes in chromatin accessibility.

To resolve between those possibilities, we quantitatively estimated the effect of *SMARCA2* and *BPTF* knockdown on glucocorticoid-dependent changes in chromatin accessibility. To do so, we fit a negative binomial linear regression model per open chromatin site with an interaction term to quantify knockdown-specific effects on chromatin accessibility. After knockdown of the glucocorticoid receptor itself, 8,588 dex-responsive regions of the genome had significantly altered chromatin accessibility **(Fig. 3C, Supp. Fig. 8A-B)**. We term those GR-dependent sites. In the BPTF knockdown samples, there were only nine significant interactions, indicating that BPTF had limited effects on glucocorticoid-dependent chromatin accessibility **(Supp. Fig. 8A,C)**. Meanwhile, in the SMARCA2 knockdown samples, we identified 606 regions of the genome that exhibited altered changes in chromatin accessibility, suggesting a substantial effect of SMARCA2 knockdown. SMARCA2 reduced the overall effect of dexamethasone on chromatin accessibility, as demonstrated by the fact that the direction of interaction effects are opposite the direction of the main effect at 97% (N = 586) of these sites. We focus on those 586 SMARCA2-dependent sites for subsequent analyses **(Fig. 3D, Supp. Fig. 8A,D).**

The effect of SMARCA2 knockdown on chromatin accessibility was not significantly different from the effect of NR3C1 knockdown at the SMARCA2-dependent sites (p = 0.79, mean of differences = 0.005, paired t-test). In contrast, at the non-SMARCA2-dependent but GR-dependent sites, *NR3C1* knockdown had a much stronger effect than *SMARCA2* knockdown (p < 2.2×10^-16^, mean of differences = 0.91, paired t-test). Together, these results show that SMARCA2 is essential for a distinct and substantial fraction of the glucocorticoid response in A549 cells, in agreement with the gene expression results above.

SMARCA2 was predominantly necessary for increasing chromatin accessibility in response to dexamethasone. Specifically, the majority of SMARCA2 knockdown interaction effects (81%) were negative effects at sites that increased chromatin accessibility with dexamethasone. For comparison, in the non-targeting samples, only 47% of regions increased accessibility in response to dex treatment. We also found that SMARCA2-dependent genomic regions exhibited larger increases in chromatin accessibility after dex treatment compared to non-SMARCA2 dependent regions ( p < 2.2×10^-16^, Welch’s T-Test). This was true even when comparing only regions with increasing chromatin accessibility ( p < 2.2×10^-16^, Welch’s T-Test) **(Supp. Fig. 9)**.

Together, these results show that SMARCA2 is necessary for dex-induced changes in chromatin accessibility at a small fraction of dex-responsive genomic regions, but that these regions are enriched for greater increases in chromatin accessibility.

### SMARCA2 dependent chromatin remodeling contributes to nearby GC-responsive gene expression

SMARCA2-dependent genomic regions were enriched near glucocorticoid-responsive genes. Specifically, 18% of the dex-responsive and GR-dependent genomic regions we identified were closest to a dex-responsive gene, while 30% of SMARCA2-dependent genomic regions were closest to a dex-responsive gene **(Fig. 3E-G)**. We hypothesized that dex-responsive genes close to genomic regions with a SMARCA2-dependent change in accessibility would exhibit larger dex-induced changes in gene expression due to the larger increases in accessibility at these SMARCA2-dependent regions compared to other GR-dependent regions. We found that to be the case. Specifically, the dex-responsive genes near regions of the genome with SMARCA2-dependent changes in chromatin accessibility do exhibit larger increases in gene expression compared to dex-responsive genes near regions of the genome with GR-dependent changes in chromatin accessibility (p = 0.00045, delta log_2_FC = 0.32, Welch’s T-Test). However, increases in gene expression were not significantly larger for dex-activated genes near genomic regions with a SMARCA2-dependent change in chromatin accessibility compared to those near a genomic region with a GR-dependent change in chromatin accessibility (p = 0.07, delta log_2_FC = 0.15). These results show that this effect is mostly due to SMARCA2-dependent regions being enriched near genes whose expression increases after dex treatment (76% of SMARCA2-dependent nearest genes vs. 55% of GR-dependent nearest genes) and not due to larger increases in expression of nearby dex-activated genes.

We also found that dex-responsive genes near SMARCA2-dependent genomic regions had more dex-responsive gene regulatory elements than genes near other GR-dependent dex-responsive regions. Glucocorticoid-responsive genes near genomic regions with SMARCA2-dependent changes in accessibility had an average of 4.9 nearby genomic regions with a dex-dependent accessibility change, while glucocorticoid-responsive genes near genomic regions with GR-dependent changes in accessibility had an average of 3.4 nearby genomic regions with a dex-dependent accessibility change (p = 3.93×10^-6^, Welch’s T-test). **(Supp. Fig 10A)**. Of those 4.9 dex-responsive regions, only ∼1 was typically SMARCA2-dependent, suggesting SMARCA2 can have very specific effects within complex gene regulatory loci **(Supp. Fig 10B)**. That regulation of a specific subset of genomic regions may explain how SMARCA2 impacts specific genes, and sometimes has weaker effects than *NR3C1* knockdown. For example, if SMARCA2 knockdown causes one enhancer element at a given locus to no longer respond to dex, but others still do, we expect that we would see smaller changes in transcription of that gene, but not for the gene to necessarily become entirely non-responsive to glucocorticoids.

### Genomic regions with a SMARCA2-dependent change in chromatin accessibility exhibit GC-induced enhancer activity

We next investigated gene regulatory activity at SMARCA2-dependent genomic regions. To do so, we used data from a genome-wide high-throughput reporter assay (STARR-seq) of dex-dependent changes in regulatory activity in the same A549 cells^3^. Importantly, those assays are episomal constructs that measure the regulatory activity of DNA sequences independent of chromatin context. Overall, SMARCA2-dependent sites had much greater dex-specific regulatory element activity than other GR-dependent regions, suggesting that SMARCA2-dependent sites have distinct DNA sequences that drive strong regulatory activity. That difference is specific to the glucocorticoid responsive regulatory activity, as demonstrated by our observation that SMARCA2-dependent and GR-dependent sites had similar regulatory activity in cells treated only with vehicle control **(Fig. 4A)**. However, in the dex condition, the SMARCA2-dependent regions of the genome had much higher regulatory activity than the GR-dependent regions or regions. That result generalized to other GR binding sites, where SMARCA2-dependent sites also had more regulatory activity than sites with GR binding according to ChIP-seq but no dex-dependent change in chromatin accessibility **(Fig. 4B)**. To quantify these differences, we compared regulatory element activity in the dex and vehicle conditions by summing the total reporter assay signal over the length-normalized regions of the genome with SMARCA2 or GR-dependent changes in accessibility **(Supp. Fig. 11)**. We found that the regions with an increase in accessibility had significantly more regulatory activity after dex treatment, while the regions with GR ChIP peaks, but no change in accessibility, did not (p < 10^-8^, Two-way ANOVA and Tukey HSD post-hoc test). SMARCA2-dependent regions also had significantly more regulatory activity after dex treatment than GR-dependent regions or regions with GR ChIP peaks (p < 10^-8^, Two-way ANOVA and Tukey HSD post-hoc test). Together, these results indicate that the DNA sequences of SMARCA2-dependent sites have greater regulatory activity within the overall glucocorticoid response.

**Figure 4.**
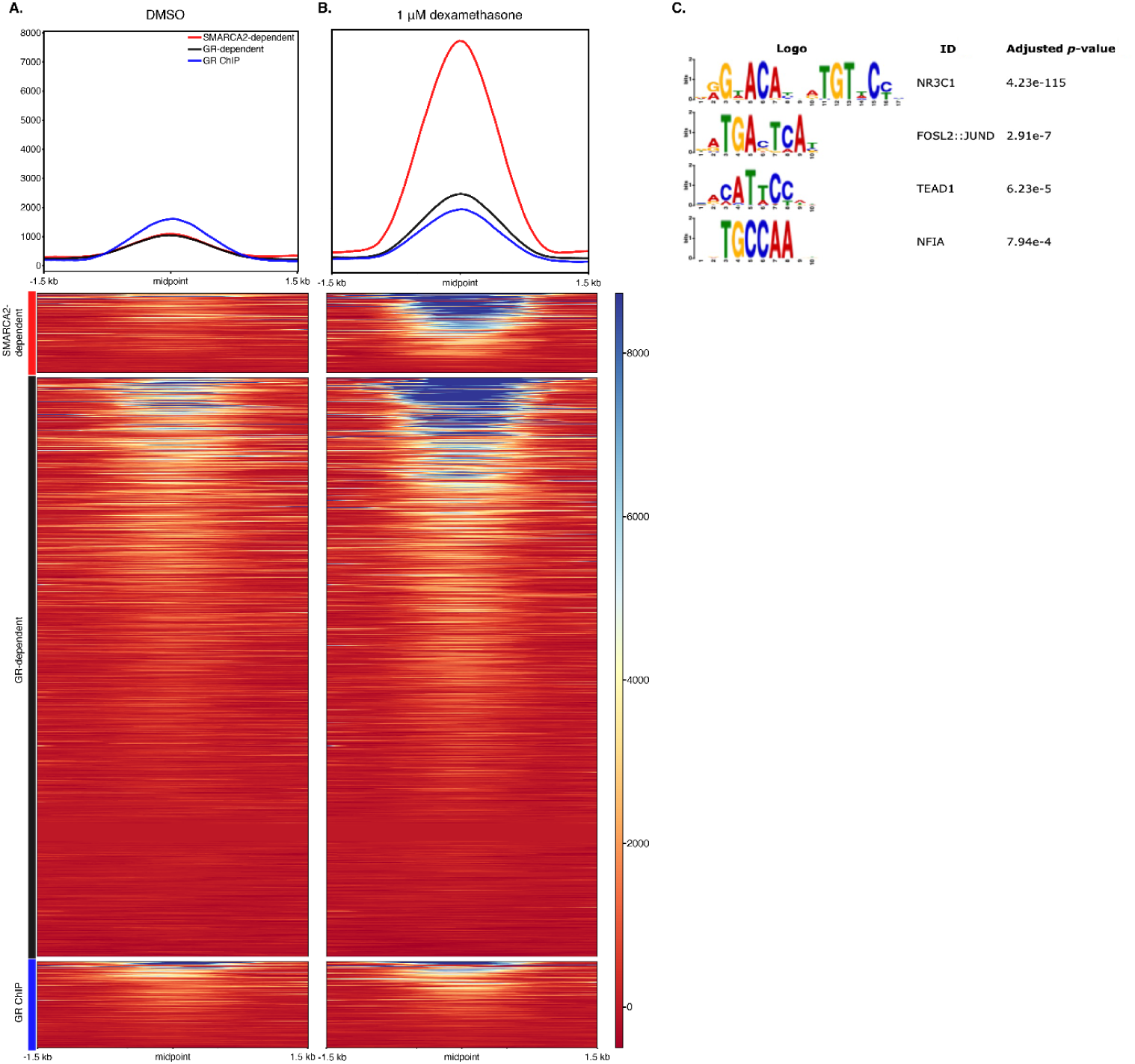
SMARCA2-dependent genomic regions exhibit enhancer activity in STARR-Seq. **A,B.** Heatmaps and profile plots showing STARR Seq activity in A549 cells treated with **A.** DMSO or **B.** 1 uM dexamethasone. The red lines show SMARCA2-dependent, dex-responsive regions, the black lines show GR-dependent, dex-responsive regions, and the blue lines show regions with GR binding but no change in accessibility. **C.** Motif enrichment using MEME2 shows TF motifs that are significantly enriched in SMARCA2-dependent genomic regions vs. all other dex-responsive genomic regions

### SMARCA2 supports increased binding of GR and other TFs to dex-responsive sites

Based on those observations, we next sought to identify a specific set of transcription factors that are enriched for binding at the SMARCA2-dependent regions of the genome. We first performed motif enrichment on the genomic regions with either a SMARCA2 or GR dependent change in chromatin accessibility. Both the SMARCA2 and GR-dependent regions of the genome with differential accessibility were enriched for factors related to the glucocorticoid response, including NR3C1, AP-1, FOXO1, and GATA2 when compared to regions of the genome without a change in chromatin accessibility after dex treatment **(Supp. Data 1)**. However, the SMARCA2-dependent regions were more enriched than GR-dependent regions for several motifs including NR3C1, AP-1, TEAD1, and NFIA motifs **(Fig. 4C)**. The NR3C1 motif had, by far, the strongest enrichment for the genomic regions with a SMARCA2-dependent change in chromatin accessibility. One possible explanation is that SMARCA2 promotes the glucocorticoid receptor binding directly to its DNA binding motif in certain contexts. The AP-1 family of transcription factors are well known to cooperate with GR to drive regulatory activity; and other studies have shown that it also synergizes with SMARCA2 to promote chromatin accessibility^40^. TEAD1 is also interesting because it is a key transcription factor in the Hippo pathway that controls cell proliferation. Previous studies have shown that TEAD family members synergize with SMARCA2 to impact cell growth in the absence of SMARCA4^40^.Together, these results indicate that specific DNA sequence motifs may dictate SMARCA2-dependent glucocorticoid responses in the genome.

As an alternative strategy to identify characteristics of SMARCA2-dependent glucocorticoid responses, we used ChIP-seq data. In a previous study, we performed ChIP-seq in A549 cells for several transcription factors and histone modifications at 0, 1, 4, and 8 hours after 100 nM dex treatment^11,41,42^, and we also completed ChIP-seq for several nuclear receptor co-factors (NCOA2, NCOA3, NCOR1, NCOR2) for this study. To understand the characteristics of the genomic regions with a SMARCA2 or GR-dependent increase in accessibility, as well as those with GR ChIP signal but no change in accessibility, we first examined the ChIP signal across these regions for all ten factors after treatment for four hours with either 100 nM dexamethasone or a 0.1% ethanol control **(Fig. 5A-B)**. In the vehicle condition, nearly all the regions with no change in accessibility showed strong AP-1 and H3K27ac signals, and were also enriched for RAD21, P300, and NCOR2 signals. These are marks of active enhancer activity and corroborate the stronger regulatory activity seen in the vehicle condition at these sites. About a third of the GR-dependent regions were similar to those with no accessibility change, while the rest were only enriched for RAD21. The SMARCA2 sites were not enriched for any factors that we tested, except for a small fraction of sites that showed AP-1 signal, which aligns with the low regulatory activity seen at these sites. However, in the dex condition, the genomic regions with no change in accessibility showed increased GR ChIP signal, but very little change in cofactor binding. The GR-dependent genomic regions showed a large increase in GR signal, as well as the recruitment of several cofactors, but most strongly at sites that already had AP-1 binding and H3K27ac in the vehicle condition. SMARCA2-dependent sites showed the strongest GR signal of any category, and saw an increase in ChIP signal for all ten cofactors, providing a potential explanation for the greatly increased regulatory signal seen in the dex condition at these sites.

**Figure 5.**
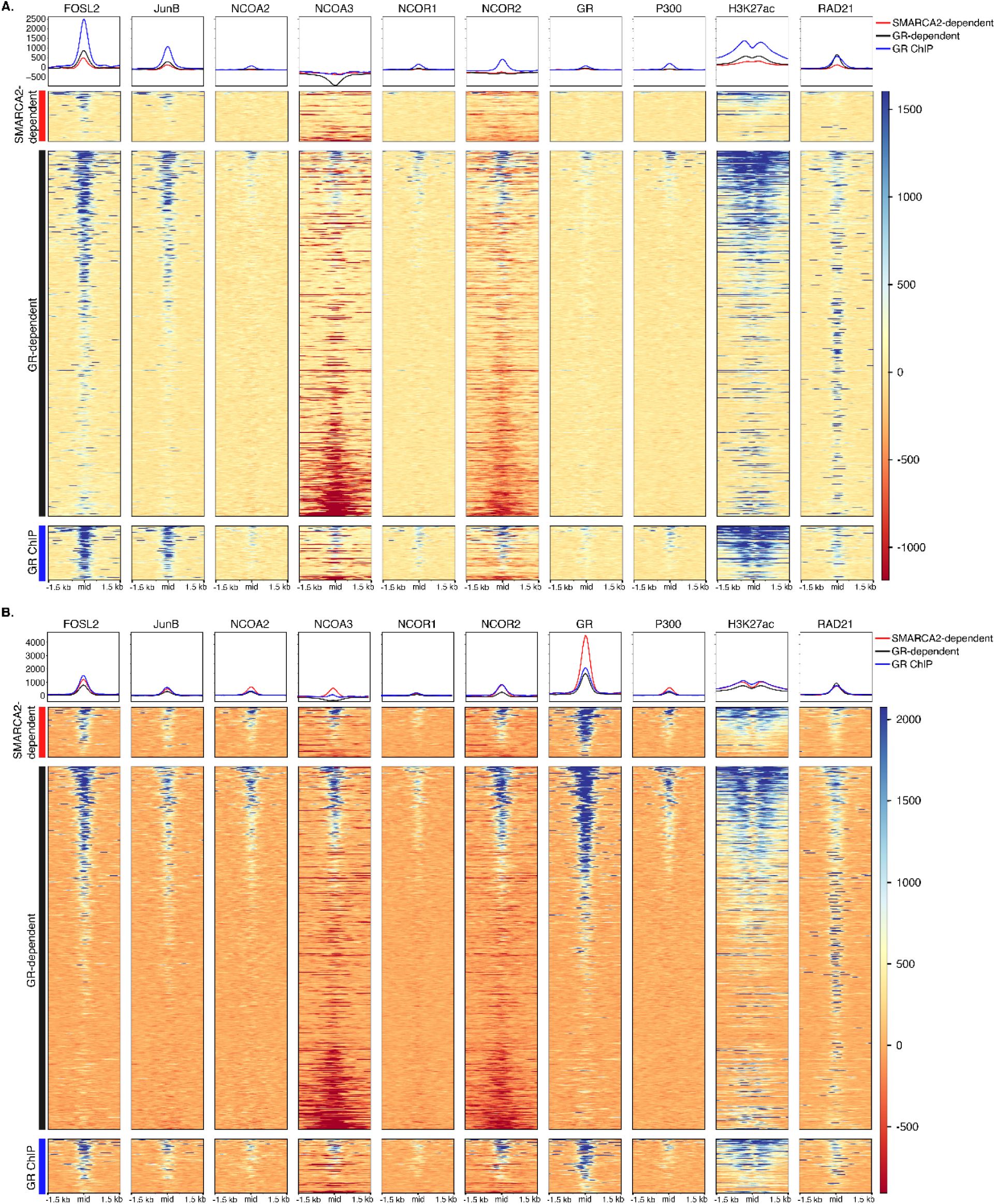
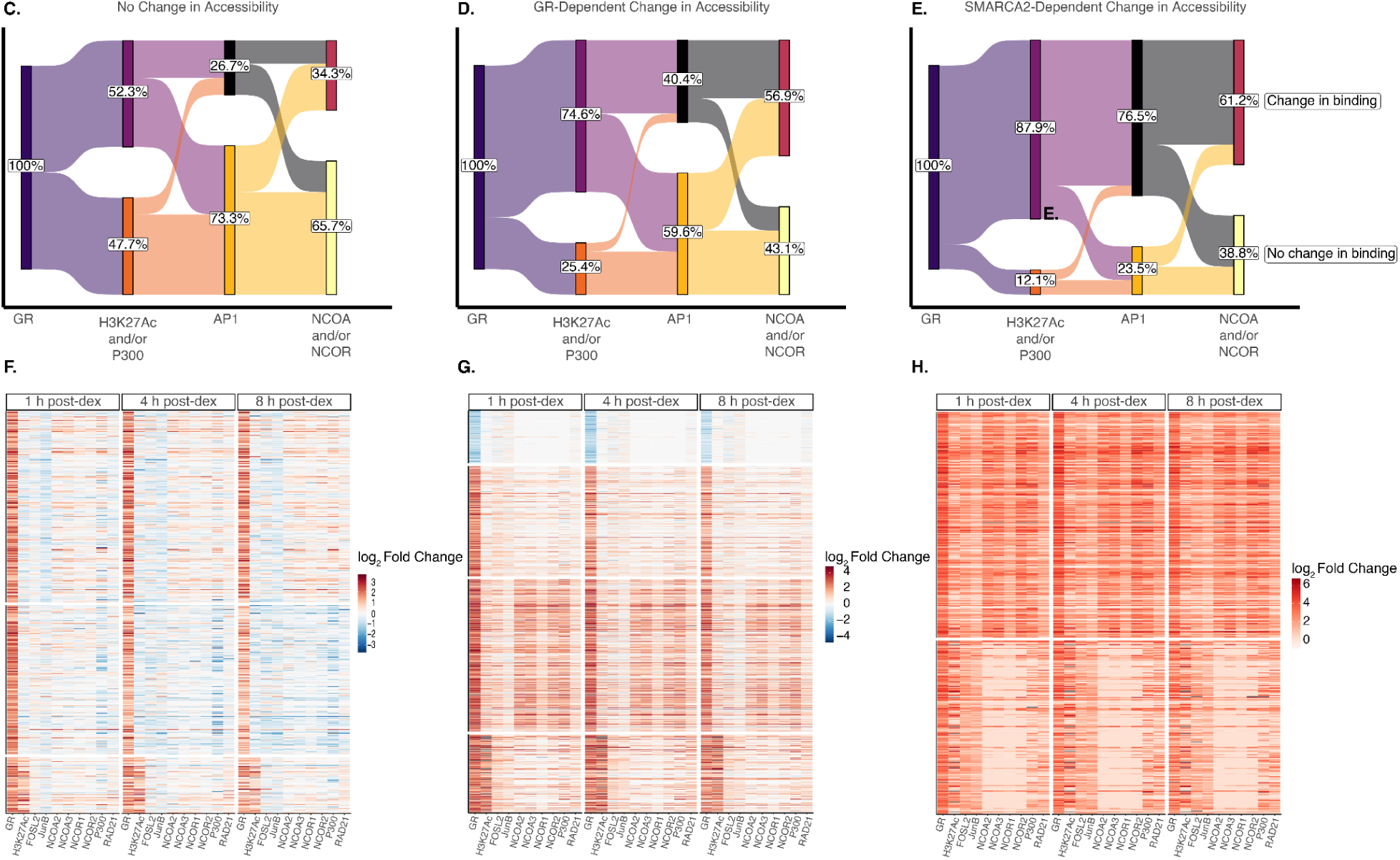
SMARCA2-dependent sites are enriched in TF binding. **A.** Sankey diagram showing the proportion of genomic regions with differential TF binding at 1, 4, or 8 hours after dex treatment for genomic regions with differential GR binding but no change in accessibility after dex treatment. **B.** Heatmap or **C.** Line plot, showing differential TF binding at 1, 4, and 8 hours post-dex treatment genomic regions with differential GR binding but no change in accessibility after dex treatment. The heatmap and line plots are divided into three different groups identified using k-means clustering. The thicker line in the line plot shows the mean log2FC of all regions in that cluster. **D-F.** The same as **A-C**., but for regions of the genome that exhibit differential GR binding as well as GR-dependent changes in accessibility after dex treatment. **G-I.**, The same as **A-C**., but for regions of the genome that exhibit differential GR binding as well as SMARCA2-dependent changes in accessibility after dex treatment.

To better understand the changes in GR and cofactor ChIP signal over time, we identified differential ChIP-seq signals (vs. time 0) from this dataset at a false discovery rate of 10% for GR, H3K27ac, FOSL2, JunB, NCOA2, NCOA3, NCOR1, NCOR2, P300, and RAD21. We then intersected these differential ChIP-seq peaks with the ATAC-seq peaks identified in our study. Since only regions of the genome that experienced an increase in GR signal seemed to experience changes in cofactor ChIP signal, we focused our analysis on regions of the genome that experienced a significant change in GR ChIP signal from baseline (time 0) to any of the 1, 4, or 8 hour post-dex timepoints (96% of genomic regions with a SMARCA2-dependent increase in chromatin accessibility and 81% of genomic regions with a GR-dependent increase in chromatin accessibility). As above, we further subset these regions into (i) those with no change in chromatin accessibility upon dex treatment; (ii) those that intersected a genomic region with a GR-dependent, non-SMARCA2 dependent increase in chromatin accessibility; and (iii) those that intersected a genomic region with a SMARCA2-dependent increase in chromatin accessibility. We hypothesized that these three subsets of genomic regions with differential GR binding upon dex treatment would have different patterns of differential ChIP-seq signal for the transcription factors and histone modifications we assayed.

The genomic regions with a SMARCA2-dependent increase in chromatin accessibility had the most dex-induced changes in ChIP-seq signal for several different transcription factors, suggesting broad recruitment of transcription factors to those sites. In contrast, the genomic regions with no change in accessibility had the least dex-induced transcription factor binding. Specifically, we determined the percentage of genomic regions for each subset that had significant differential ChIP signal for H3K27ac and/or P300, AP-1 factors (FOSL2 or JunB), or the nuclear receptor coactivators (NCOA2, NCOA3) and corepressors (NCOR1, NCOR2) at any time point **(Fig. 5C-E)**. The genomic regions with a SMARCA2-dependent increase in chromatin accessibility were most enriched for differential dex-induced ChIP-seq signals across all of these categories, but had the largest difference for the AP-1 factors. Most (77%) genomic regions with a SMARCA2-dependent increase in chromatin accessibility exhibited dex-induced differential AP-1 ChIP signal, compared to 40% of genomic regions with a GR-dependent increase in chromatin accessibility and 27% of genomic regions with no change in accessibility. This result agrees with the fact that AP-1 motifs are enriched in genomic regions with a SMARCA2-dependent change in chromatin accessibility compared to those with a GR-dependent change in accessibility. Additionally, regions of the genome with both GR- and SMARCA2-dependent dex-induced increases in accessibility had a greater percentage of regions with differential ChIP activity than the regions with no dex-induced change in accessibility across all categories **(Fig. 5C-E)**. This difference became more stark when quantifying the proportion of genomic regions with increased ChIP signal after dex treatment **(Supp. Fig. 12)**. This difference in ChIP signal could not be explained by differences in the length or baseline accessibility of the genomic regions with a SMARCA2-dependent change in accessibility, GR-dependent change in accessibility or no change in accessibility **(Supp. Fig. 13)**. This result agrees with previous work which found a set of genomic regions that showed GR binding, but no dex-dependent increase in regulatory activity or TF binding except GR^16^. These regions had high AP-1, EP300, and H3K27ac ChIP signals in the vehicle condition, and it is suspected that the GR binds to these regions via AP-1 as opposed to direct DNA binding^7,16^. These results demonstrate that the genomic regions with increased accessibility after dex treatment mostly experience increased transcription factor binding after dex treatment, while the genomic regions with no change in accessibility after dex treatment experience a greater mix of increasing and decreasing transcription factor binding after dex treatment.

Interestingly, SMARCA2-dependent regions were the only one of the three subsets that was not enriched for RAD21 ChIP-seq signal at baseline. Previous work has shown that glucocorticoid-responsive chromatin interactions are mostly mediated through more frequent interactions of pre-existing chromatin contacts as opposed to new loop formation, and that these contacts are depleted in CTCF but enriched for RAD21^41^. The GR-dependent regions of the genome showed RAD21 enrichment in the vehicle condition even at sites without AP-1 binding or H3K27ac, but the SMARCA2-dependent sites did not. This suggests that SMARCA2-dependent sites may also represent sites of new loop formation, or, possibly, that due to the large numbers of TFs at these sites, RAD21 is acting as a scaffold to stabilize those interactions^43,44^.

Finally, we investigated whether patterns of transcription factor binding over time differ depending on the impact of SMARCA2 **(Fig. 5F-H)**. Overall, SMARCA2-dependent sites **(Fig 5H)** had much greater dex-responsive increases in the binding of all transcription factors and co-factors measured across all time points. Indeed, more than half of the SMARCA2-dependent sites had substantial and persistent binding of nearly all other factors, cofactors, and activating histone modifications. GR-dependent sites had a more moderate overall increase in transcription factor and co-factor binding. A smaller fraction of those sites showed strong recruitment of all other factors. The GR-dependent sites also included a group of repressed sites not found in the SMARCA2 dependent group. Finally, sites bound by GR but without differential chromatin accessibility recruited other transcription factors and co-factors the least. Together, these results suggest that SMARCA2 contributes to GR responses by increasing GR and potentially other transcription factors binding to their cognate motifs and, in turn, broadly increasing recruitment of activating gene regulatory factors to those sites.

## Discussion

Genome-wide knockout screens using CRISPR-Cas9 have been powerful unbiased approaches for determining which genes are necessary for a phenotype of interest^45^. Here, we use a CRISPR knockout library targeting 19,114 genes coupled with GILZ protein staining and cell sorting to determine which genes are necessary for dexamethasone-induced GILZ expression. We identify members of two different chromatin remodeling complexes, SMARCA2, a member of the SWI/SNF complex, and BPTF, a member of the NURF complex, from this screen.

NURF has not previously been shown to have a role in the glucocorticoid response, but has been shown to be a co-activator of a fly nuclear receptor.^46^ NURF appears to have a very specific role in the glucocorticoid response – BPTF only affected the expression of ∼24 genes and the chromatin accessibility of nine genomic regions at an FDR of 10%. It is possible that the complex does play a larger role, but because BPTF knockdown was not as strong as knockdown of other target genes (70% vs. 83% for SMARCA and > 90% for NR3C1), this small amount of BPTF was sufficient for chromatin remodeling activity. Regardless, perturbation of BPTF is a way to modify a very specific subset of the glucocorticoid response, and the mechanism by which it affects target gene expression is an area for further study.

The SWI/SNF complex has been well-characterized as having a role in the glucocorticoid response, but that work has largely focused on the role of SMARCA4. Furthermore, previous studies of SMARCA2’s role in the glucocorticoid response have focused on synthetic reporters or a handful of highly glucocorticoid-responsive genes^17,18,25–27^. Here, we expand upon those studies by using genome-wide, unbiased methods to characterize the role of SMARCA2 across the entire glucocorticoid response. Similarly to previous results with SMARCA4, SMARCA2 is responsible for gene expression changes for a substantial portion of the glucocorticoid response (27%), and can affect the expression of both dex-activated and dex-repressed genes, though it impacts more dex-activated genes^30^. SMARCA4 is commonly mutated in cancer, including in A549 cells, and so the result that SMARCA2 can still facilitate the glucocorticoid response is important, since SMARCA2 and SMARCA4 can only sometimes compensate for the role of the other^6,47,48^.

Interestingly, other previous work has raised the question of redundancy in glucocorticoid-induced chromatin accessibility changes beyond just the SMARCA2/SMARCA4 pair. One study showed that, in yeast, overexpression of the Ada2 protein (human homologs TADA2A and TADA2B) could partially rescue glucocorticoid receptor activity in a SWI/SNF deficient yeast strain^27^. Another study of SMARCA2 in A549 cells performed ChIP-qPCR for SMARCA2 at a handful of genomic regions with glucocorticoid-induced changes in accessibility^17^. They found that SMARCA2 ChIP signal increased similarly after dex treatment at loci with SMARCA2-dependent changes in gene expression as well as those that were SMARCA2-independent^17^. One possible explanation given by the authors for the fact that SMARCA2 ChIP signal increased at certain genomic regions even though SMARCA2 knockdown had no impact on gene expression or chromatin accessibility at that region, is that there is redundancy at those loci with other chromatin remodeling complexes. This certainly seems possible, especially given that knockdown studies show SWI/SNF is not responsible for chromatin remodeling at all glucocorticoid-responsive regions. That, coupled with the fact that we did not identify other chromatin remodeling complexes in our genome-wide screen, except for NURF as explained above, suggests that either only SWI/SNF is necessary for chromatin remodeling at the TSC22D3 locus, or there is redundancy with other chromatin remodeling complexes.

The subset of genomic regions we identified to have SMARCA2-dependent increases in chromatin accessibility also have many hallmarks of dex-induced enhancer activity. Those include very high dex-specific regulatory activity, increased binding of many GR co-factors after dex treatment, and strong enrichment of the GR DNA binding motif and GR ChIP signal, even compared to other dex-responsive sites. Previous work has shown that only a fraction of GR ChIP signal is a result of direct GR binding, but that GR binding sites are characterized by lower baseline accessibility, higher changes in chromatin accessibility after dex treatment, and higher dex-induced regulatory activity in STARR-seq^16^. It has also been shown that the GR motif and change in GR binding at an enhancer are strong predictors of P300 binding, another mark of active enhancers^11,16^. SMARCA2-dependent regions match all of these criteria, and previous work shows a similar role for SMARCA4^7,31^. In fact, one of those studies showed that a GR mutant unable to bind to regions of the genome with SMARCA4-dependent increases in accessibility showed an 80% reduction in significantly dex-responsive genes despite these sites only comprising ∼25% of GR binding sites, suggesting they have a critical regulatory function^31^.

## Acknowledgements

Funding for this project was provided by NIH grants (RM1HG011123, UM1HG009428, and U01HG007900 to TER and CAG). We thank Nicholas S. Giroux for guidance on integrating ATAC-seq data with STARR-seq and ChIP-seq data.

## Author Information

## Contributions

T.E.R. and C.A.G. conceived, funded, and supervised the study. D.D.K. performed the genome-wide CRISPR screen and gRNA validation. S.M.M. contributed to the siRNA knockdown validation of SMARCA2 and BPTF, performed the RNA-seq and ATAC-seq experiments, and analyzed the resulting data. K.S. contributed to the ATAC-seq study. C.W. and L.C.B performed ChIP-seq. A.G.J. contributed to the siRNA knockdown validation of SMARCA and BPTF. A.B. and R.V. created the pre-processing pipelines for RNA-seq, ATAC-seq, and ChIP-seq data and contributed to the bioinformatic analyses in this paper. G.D.J. performed the STARR-seq experiments and contributed to integrated analyses of ATAC-seq and STARR-seq data. T.E.R. and S.M.M. wrote the manuscript with input from co-authors.

## Methods

### Cell Culture

The A549 cell line was obtained from ATCC. It was authenticated using STR profiling through the Duke Cell Culture and DNA Analysis facility. Mycoplasma testing was performed by Eurofins. A549 cells were grown in humidified incubators at 37 °C and 5% CO_2_. Cells were cultured using F12K (Gibco) + 10% FBS (Gibco) + 1% penicillin-streptomycin (Gibco)

### Genome-wide CRISPR screen

To conduct a genome-wide knockout CRISPR screen, we used the one-vector version of the 77,441 member Brunello Human CRISPR knockout pooled library (Addgene 73179) which targets 19,114 genes^12^. A549 cells were transduced at low MOI at 300x coverage. After three days, cells were selected with 1 ug/mL puromycin (Gibco). After passaging for eight days, cells were treated with 100 nM dexamethasone or an ethanol vehicle control for 4 hours. We performed five replicates of this screen.

After drug treatment, cells were fixed and stained with an anti-GILZ antibody (eBioscience Ref. 14-4033-82, clone CFMKG15) at a 1:1000 dilution using the eBioscience Foxp3 / Transcription Factor Staining Buffer Set (Invitrogen). The cells were then resuspended in flow cytometry staining buffer (eBioscience) and sorted using a SONY SH800 sorter. Cells were first gated on single cells using FSC/SSC. Next cells were gated on PE in the FL2 channel, which marked GILZ expression. The top and bottom 10% of PE-labelled cells were then sorted into high and low GILZ bins.

Genomic DNA was extracted from bulk-transduced cells as well as those from cells sorted into high and low GILZ expression bins using the FFPE miniprep kit (Zymo).

### Genomic DNA Sequencing

To amplify the gRNA libraries from each sample, 8.3 μg of gDNA was used as template across 8 100 μL PCR reactions using Q5 hot start polymerase (NEB, M0493L). Amplification was carried out following the manufacturer’s instructions using 25 cycles at an annealing temperature of 60 °C using the following primers:

Fwd 5′-AATGATACGGCGACCACCGAGATCTACACAATTTCTTGGGTAGTTTGCAGTT

Rev 5′-CAAGCAGAAGACGGCATACGAGAT GACTCGGTGCCACTTTTTCAA

Amplified libraries were purified using Agencourt AMPure XP beads (Beckman Coulter, A63881) using double size selection of 0.65X and then to 1X the original volume. Each sample was quantified after purification using the Qubit dsDNA High Sensitivity Assay Kit (ThermoFisher, Q32854). Samples were pooled and sequenced on a MiSeq (Illumina) with 21bp paired-end sequencing using the following custom read and index primers:

Read1 5′-GATTTCTTGGCTTTATATATCTTGTGGAAAGGACGAAACACCG

Index 5′-GCTAGTCCGTTATCAACTTGAAAAAGTGGCACCGAGTC Read2 5′-GTTGATAACGGACTAGCCTTATTTTAACTTGCTATTTCTAGCTCTAAAAC

### Data Processing and Enrichment Analysis

FASTQ files were aligned to custom indexes (generated from the bowtie2-build function) using Bowtie255 with the options -p 32 --end-to-end --very-sensitive -3 1 -I 0 -X 200. Counts for each gRNA were extracted and used for further analysis. All enrichment analysis was performed using R. For DHS level analysis, gRNAs for each DHS were grouped together and a linear regression model (normalized_gRNA_count = β1*(sorted_bin)+ β2*(replicate)) was used to detect differences between the high and low conditions using the Holm method for multiple hypothesis correction. For individual gRNA enrichment analysis, the DESeq256 package was used to compare between high and low, unsorted and low, or unsorted and high conditions for each screen.

### Analysis of CRISPR Screen Data

FASTQ files were aligned to custom indexes (generated from the bowtie2-build function) using Bowtie255 with the options -p 32 --end-to-end --very-sensitive -3 1 -I 0 -X 200. Counts for each gRNA were extracted and used for further analysis. Raw read counts can be found in **Supp. Data 2** MaGeck was run on the un-normalized read counts using the mageck test command. Control normalization to the 1,000 non-targeting gRNAs in the library was used. High and low bins for each drug condition were analyzed separately. To create a single log_2_ fold change for each gene, we averaged the high and low bin results as follows: (High_bin_log_2_FC - Low_bin_log_2_FC)/2. To calculate a combined log_10_ RRA score, if gene behavior was concordant between the two bins (i.e. depleted from the high bin and enriched in the low bin or vice versa), we added the log_10_(RRA Score) for the high and low bins, while if gene behavior was discordant (i.e. enriched or depleted from both bins), we subtracted the log_10_(RRA Score) for the high and low bins.

### Followup Screen Validation

The protospacers from the top enriched gRNAs found in the screen were ordered as oligonucleotides from IDT and cloned into a lentiviral gRNA expression vector (lentiCRISPR v2, Addgene Plasmid #52961). Lentivirus for each gRNA was produced in a 96-well format in HEK cells and the resulting supernatant was used to infect A549 cells in replicate transductions. 3 days post infection, A549 cells were selected using puromycin and grown for 5 more days before cells were harvested for analysis by flow cytometry and RT-qPCR to assay for GILZ protein levels and RNA levels, respectively.

RT-qPCR was performed using the Cells to Ct (Invitrogen, A57985) with the FX96 Real-Time PCR Detection System (Bio-Rad) using multiplex Taqman probes for TSC22D3 (Hs00608272_m1) and GAPDH(4326317E). The results are expressed as fold-increase mRNA expression of the gene of interest normalized to GAPDH expression by the ΔΔCt method. For flow cytometry analysis, staining of cells was conducted using the GILZ antibody as described below and analyzed using the MACSQuant VYB flow cytometer (Miltenyi Biotec).

### siRNA Validation

#### Transfection

A549 cells were seeded at a density of 20,000 cells/cm^2^ in F12K + 10% FBS + 0.25% penicillin-streptomycin. The next day, cells were transfected with ON-TARGET siRNA SMART pools (Horizon Discovery) targeting genes of interest using DharmaFECT 1 transfection reagent (Horizon Discovery) according to manufacturers protocols. We used 1 uL of DharmaFECT 1 per well of a 24-well plate and 4uL of DharmaFECT 1 per well of a 6-well plate.

#### GILZ Staining and Flow Cytometry

A549 cells transfected with siRNAs were treated with 100 nM dexamethasone or 0.1% ethanol as a vehicle control 48 hours post-transfection (n = 4 independent transfections). 24 hours after drug treatment, cells were fixed and stained with an anti-GILZ antibody (eBioscience Ref. 14-4033-82, clone CFMKG15) using the eBioscience Foxp3 / Transcription Factor Staining Buffer Set (Invitrogen) according to manufacturers protocols. Fixation was performed for 40 min, blocking with 2% normal mouse serum was performed for 15 min, and anti-GILZ antibody staining was performed for 35 min at a 1:1000 dilution. After staining, cells were resuspended in 150 uL of flow cytometry staining buffer (eBioscience) and analyzed on an Accuri C6 plus flow cytometer (BD). First, cells were gated on FSC and SSC to determine live cells and single cells. Next, FITC MFI was determined for all samples using a gated cutoff of 15,000 single cells.

#### RNA Extraction and RT-qPCR

A549 cells transfected with siRNAs were treated with 100 nM dexamethasone or 0.1% ethanol as a vehicle control 48 hours post-transfection (n = 3 independent transfections). Two hours after drug treatment, RNA was harvested from cells using the RNeasy mini kit (Qiagen) and cDNA was synthesized using the high-capacity cDNA reverse transcription kit (Applied Biosystems). Multiplexed qPCR was performed using TaqMan Gene Expression Master Mix (Applied Biosystems), a FAM-labeled Taqman assay (Thermo) for the appropriate target gene, and a VIC-labeled endogenous control assay (Thermo) for GAPDH. qPCR was performed on a StepOne Plus thermal cycler under the following conditions:

95 °C for 10 min

40x

95 °C for 15 seconds

60 °C for 1 minute

### Generation of FLAG tagged cell lines

A549 cells were electroporated in a 0.2-cm cuvette using Bio-Rad’s GenePulser Xcell; 2 × 10^6^ cells were resuspended in 200 µL of Opti-MEM (Thermo Fisher Scientific) with 5 µg of pooled plasmid (1 µg of donor, 1 µg of guide RNA, and 3 µg of Cas expression vectors). This was done in biological triplicate for each guide RNA used, where three separate electroporations were performed for each guide RNA and these were maintained independently thereafter. Immediately after electroporation, cells were rescued with 1 mL of complete media and transferred into complete media. Media was exchanged every 2 d thereafter. Transfection efficiencies were routinely higher than 60%, as determined by fluorescence microscopy after delivery of a control eGFP expression plasmid. Five days postelectroporation, cells were selected with puromycin at a concentration of 1 µg/mL. Selection was performed for 3 d. Cells were then treated with adenoviral Cre recombinase per the manufacturer’s protocol (University of Iowa Vector Core). The efficiency of the desired editing event was assessed by restriction fragment length polymorphism analysis. Genomic DNA of the cell population was extracted using the DNeasy kit (Qiagen). The C terminus of the gene of interest was amplified using AccuPrime Taq DNA Polymerase (Thermo Fisher Scientific) with primers that bind external to the homology arm region. The PCR product was purified using a 0.5× SPRI bead purification (Beckman Coulter), and the product was digested using PsiI (New England Biolabs), which recognizes a DNA sequence embedded in the 3xFLAG coding sequence. Digests were run on an agarose gel and the efficiency of gene editing was assessed by densitometry. The desired modification was then further confirmed by western blot using the Sigma-Aldrich M2 antibody (F1804). Cells were then expanded and stored in 1 × 10^7^ cell aliquots for all subsequent experiments.

### RNA Sequencing

Using the same samples as the RT-qPCR, 1 µg of RNA per sample was prepared for RNA sequencing using the TruSeq mRNA stranded library prep kit (Illumina) according to manufacturers instructions. Libraries were quantified using a qPCR standard curve (KAPA ABI Prism qPCR Master Mix). Libraries were run using a 50 bp paired-end (100 cycle) NextSeq 2000 P3 kit at a concentration of 750 pM.

### RNA Sequencing Data Processing

RNA Sequencing data was first pre-processed using our paired end, reverse-stranded, CWL processing pipeline for RNA-seq with a known splice junction database (sjdb), which can be found at: https://github.com/ReddyLab/IGVF-cwl/l/tree/master/v1.0/RNA-seq_pipeline. All RNA-seq samples were first examined for consistent quality using FastQC v0.11.2 (Babraham Institute). Raw reads were trimmed to remove adapters and bases with average quality score (*Q*) (Phred33) of < 20 using a 4bp sliding window (SLIDINGWINDOW:4:20) with Trimmomatic v0.32 (Bolger et al., 2014). Trimmed reads were then aligned to the primary assembly of the GRCh38 human genome using STAR v2.4.1a (Dobin et al., 2013), using a A549-specific non-canonical splice junction database to maximize mapping rates. Aligned reads were assigned to genes in the GENCODE v22 comprehensive gene annotation using the featureCounts command in the subread package with default settings (v1.4.6-p4).

Next, DESeq2 was used to detect differentially expressed genes after dex treatment using the RSEM files generated by the pre-processing pipeline. We filtered out genes with low counts by requiring that each gene have a total read count across all N = 35 samples of at least 350 (an average of 10 reads per sample). The model used was:

`∼ siRNA + Drug_Condition + siRNA:Drug_Condition`

We estimated false discovery rates using the Benamini-Hochberg method as implemented in DESeq2.

### ATAC Sequencing

A549 cells transfected with siRNAs in 24 well plates were treated with 100 nM dexamethasone or 0.1% ethanol as a vehicle control 48 hours post-transfection (n = 3 independent transfections). Two hours after drug treatment, 50,000 cells per sample were harvested and processed using the Omni-ATAC protocol^49^. The transposase used was from the Tagment DNA TDE1 Enzyme and Buffer Kits (Illumina). Libraries were quantified using high sensitivity dsDNA Quant-It reagents (Invitrogen). Libraries were run at the Duke University Sequencing and Genomic Technologies Shared Resource using a 50 bp paired end (100 cycle) Nova Seq 6000 S1 kit.

### ATAC Sequencing Data Processing

ATAC sequencing data was first pre-processed using our paired end processing pipeline for paired end ATAC-seq with blacklist removal, which can be found at: https://github.com/Duke-GCB/GGR-cwl/blob/master/v1.0/ATAC-seq_pipeline/pipeline-pe-blacklis t-removal.cwl. Sequencing data quality was assessed with FastQC and adapters were trimmed from the reads with Trimmomatic. Trimmed reads were aligned to the GRCh38 genome using Bowtie (v1.0.0), reporting only alignments having no more than two mismatches, discarding multi-mapping reads and (-v 2 --best --strata -m 1). Reads mapping to the ENCODE hg38 blacklisted regions (https://www.encodeproject.org/files/ENCFF356LFX; manually curated regions with anomalous signal across multiple genomic assays and cell types) were removed using bedtools2 intersect (v2.25.0). Properly paired reads were then filtered to exclude PCR duplicates using Picard MarkDuplicates (v1.130; http://broadinstitute.github.io/picard/). Reads pileups were computed to generate counts per million (CPM) bigWig files for visualization using deeptools bamCoverage (v3.0.1).

Next, DESeq2 was used to detect differentially accessible regions of the genome after dex treatment using the count tables generated by featureCounts as part of the pre-processing pipeline. We filtered out regions of the genome with low counts by requiring that the total number of read counts for each genomic region across all N = 35 samples was at least 1,000. The model used was:

`∼ siRNA + Drug_Condition + siRNA:Drug_Condition`

We estimated false discovery rates using the Benamini-Hochberg method as implemented in DESeq2.

### ChIP Sequencing

Generation of 3x FLAG tag knock-in A549 cell lines for NCOA2, NCOA3, NCOR1, and NCOR2 and subsequent ChIP-seq were performed using the same methods as in Ref. 42.

Briefly, we first introduced C-terminal fusions of 3x FLAG epitope to target genes using homology-directed repair. We co-electroporated A549 cells with a plasmid expressing a CRISPR enzyme (Cas9 or Cas12a) to induce a double stranded break at the C-terminus of the ORF, a gRNA plasmid, and a donor template with 3x FLAG sequence to be used for double-stranded break repair. Efficiency of these edits was confirmed through restriction fragment length polymorphism analysis and western blotting.

To generate ChIP libraries, cells were treated with 100 nM dexamethasone for 0, 1, 4, or 8 hours with the timing staggered so that all cells were harvested at the same time. Each cell line was assayed in triplicate for each time point, with ∼60 x 10^6^ cells per replicate. Cells were crosslinked for 10 min at room temperature with 1% formaldehyde in media before being quenched with 0.125M glycine for 5 min at room temperature. Cells were then lysed in Farnham lysis buffer with protease inhibitor and manually scraped from the plate. Subsequently, cells were sheared in RIPA buffer with protease inhibitor using the Bioruptor Twin. ChIP was performed as in Ref. 13, reverse-crosslinked DNA was cleaned using the Qiagen PCR purification kit, and 7 ng of this cleaned DNA was input into the Kapa Biosystems Hyper Prep kit for Illumina sequencing to make sequencing libraries.

### ChIP Sequencing Data Processing

ChIP sequencing data was first pre-processed using https://github.com/Duke-GCB/GGR-cwl/blob/master/v1.0/ChIP-seq_pipeline/pipeline-se-with-control.cwl. Briefly, adapter sequences were removed from the raw reads using Trimmomatic. Reads were aligned using Bowtie v1.0.0, reporting the best alignment with up to 2 mismatches (parameters --best --strata -v 2). Duplicates were marked using Picard MarkDuplicates v1.130 (http://broadinstitute.github.io/picard/), while low mappability or blacklisted regions identified by the ENCODE project were filtered out from the final BAM files. Input subtracted signal files were generated with deeptools bamCoverage (v3.0.1) ignoring duplicates, extending reads 200bp and applying RPKM normalization. Using the sequenced input controls, binding regions were identified using the callpeak function in MACS2 v2.1.1.20160309.

Next, DESeq2 was used to detect differential transcription factor binding and histone modifications using the count tables generated by featureCounts as part of the pre-processing pipeline. We filtered out regions of the genome with low counts by requiring that the total number of read counts for each genomic region across all samples was at least 10. The model used was:

`∼ Timepoint`

We estimated false discovery rates using the Benamini-Hochberg method as implemented in DESeq2. We analyzed the AP-1 factor and GR ChIP data jointly, the NCOA and NCOR factor ChIP data jointly, and the RAD21, P300, and H3K27ac ChIP data individually.

### Transcription Factor Motif Enrichment Analysis

To perform motif enrichment, we first converted ATAC-seq regions of interest to fasta format using bedtools. The command used was:

`bedtools getfasta -fi hg38.fa -bed ATAC_peaks.bed -fo ATAC_peaks.fa`

Next, we used the AME software from MEME Suite version 4.12.0 to perform motif enrichment using the JASPAR_2018_matrix_clustering_vertebrates_central_motifs.meme database as a reference. As part of this analysis, we compared GR- or SMARCA2-dependent genomic regions to a negative control peak set with no dex-induced change in chromatin accessibility. For the GR analysis, we defined this negative set as ATAC peaks that had an absolute interaction log_2_ fold change of < 0.3 and FDR > 0.7 in the cells treated with the NR3C1-targeting siRNA. For the SMARCA2 analysis, we defined this negative set as ATAC peaks that had an absolute interaction log_2_ fold change of < 0.1 and FDR > 0.9 in the cells treated with the SMARCA2-targeting siRNA. The command used to perform motif enrichment was:

`ame --method fisher peaks_of_interest.fa --control ctrl_peaks.fa JASPAR_2018_matrix_clustering_vertebrates_central_motifs.meme`

### Integration of ATAC-seq With Other Datasets

To identify the closest gene to each genomic region identified in the ATAC-seq study, we used bedtools to identify the closest protein coding TSS from Gencode V28. The command used was:

`bedtools closest -a sorted.ATAC.peaks.bed -b Gencode.hg38.protein_coding.TSS.v28.sorted.bed -D ref`

To intersect ATAC and ChIP-seq peaks, we used bedtools to identify ATAC peaks that overlapped ChIP peaks. The command used was:

`bedtools intersect -loj -sorted -a sorted_chip_peaks.bed -b sorted.ATAC.peaks.bed`

To visualize Chip-seq signal across ATAC-seq peaks, we first used the GenomicRanges package in R to resize all the ATAC peaks to be 1 kb total length, centered on the midpoint of the original ATAC peak. We then used the computeMatrix function from deepTools followed by the plotHeatmap function to visualize ChIP-seq signal across these regions. We used mean, RPKM-normalized, input subtracted, ChIP-seq signal across two to three replicates per drug condition.

To visualize STARR-Seq signal, we again used the computeMatrix function from deepTools followed by the plotHeatmap function to visualize STARR-seq signal across these regions. To quantify STARR-Seq signal across these regions, we used the multiBigwigSummary command from deepTools to quantify the total STARR-seq signal across the resized, length-normalized ATAC peaks. This generated a text file containing the coordinates of each ATAC peak and the value of the total BigWig signal across that region. The BigWig files represented the mean, RPKM-normalized, input subtracted, STARR-seq signal across five replicates in order to allow for comparisons between the dexamethasone and vehicle treatment conditions. The command used was:

`multiBigwigSummary BED-file -b Mean_Dex_RPKM_Input_subtracted.bw Mean_Veh_RPKM_Input_subtracted.bw --BED ATAC.peaks.bed --outRawCounts`

## Data Availability

Supplementary data has been made available with this study. Raw sequence data and associated processed files will be deposited in NCBI’s Gene Expression Omnibus (Edgar et al., 2002)

## Code Availability

CWL pre-processing pipelines for RNA-seq, ATAC-seq, and ChIP-seq are available at https://github.com/Duke-GCB/GGR-cwl/tree/master/v1.0

