## Supplementary Information for "SMARCA2 is an essential and potent cofactor for a specific subset of the glucocorticoid response in A549 cells"

#### Supplementary Figures

##### Supplementary Figure 1

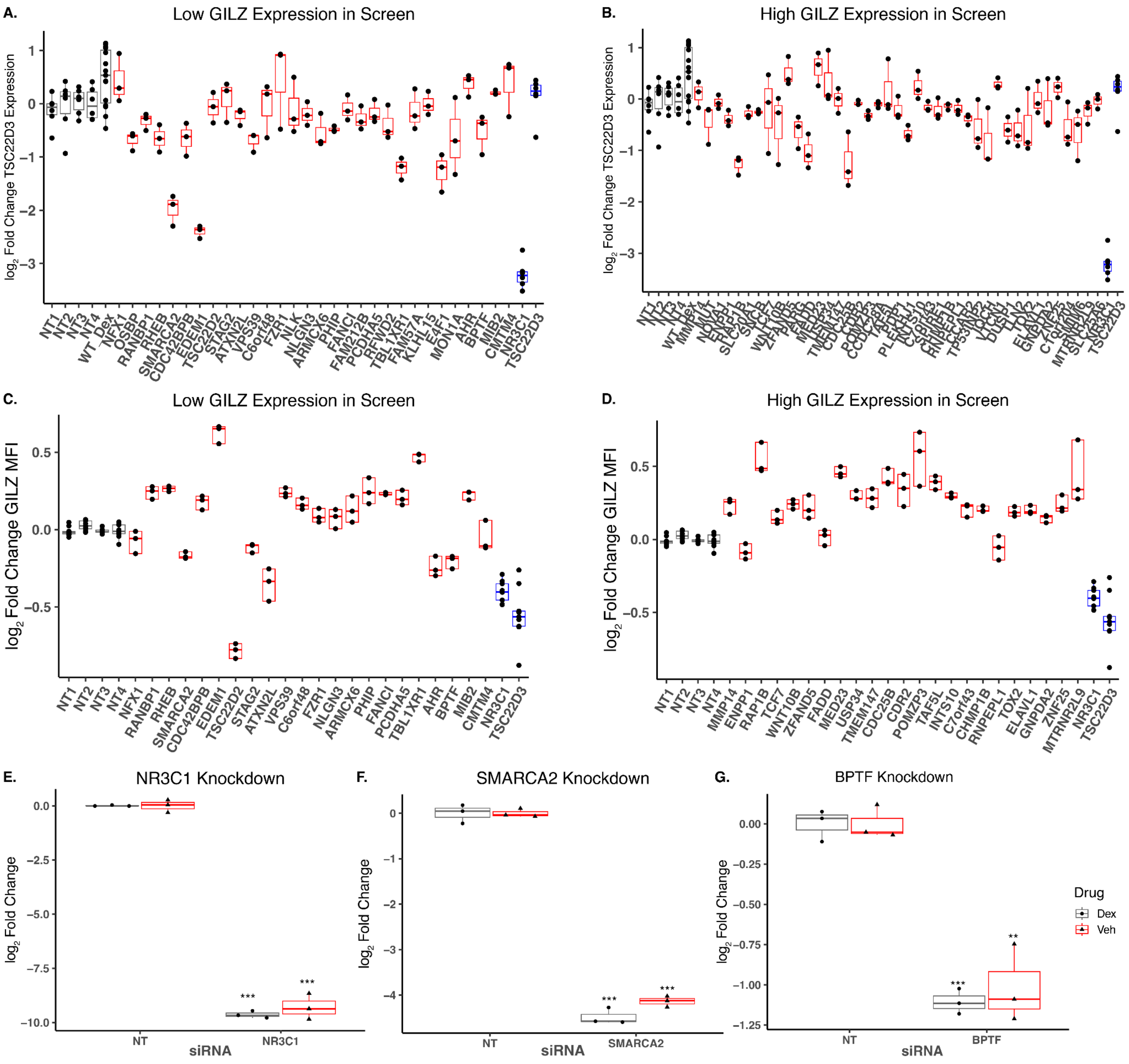


Validation of the effects of several gRNAs from the screen on TSC22D3 expression using RT-qPCR for **A.** gRNAs negative effects on glucocorticoid-induced GILZ expression in the screen and **B.** gRNAs with positive effects on glucocorticoid-induced GILZ expression in the screen (n = 3 to n = 5 independent transductions). **C,D.** Validation of the effects of several gRNAs from the screen on GILZ expression using protein staining and flow cytometry for **C.** gRNAs negative effects on glucocorticoid-induced GILZ expression in the screen and **D.** gRNAs with positive effects on glucocorticoid-induced GILZ expression in the screen. Samples transduced with non-targeting control gRNAs are shown in gray, positive control gRNAs targeting *NR3C1* and *TSC22D3* are in blue, and gRNAs to be validated in red. **E-G.** Validation of siRNA knockdown of target genes for **E.** NR3C1, **F.** SMARCA2, and **G.** BPTF using RT-qPCR (n = 3 independent transfections). * $p < 0.05$, ** $p < 0.01$, *** $p < 0.001$, one-tailed student's t-test

#### Supplementary Figure 2

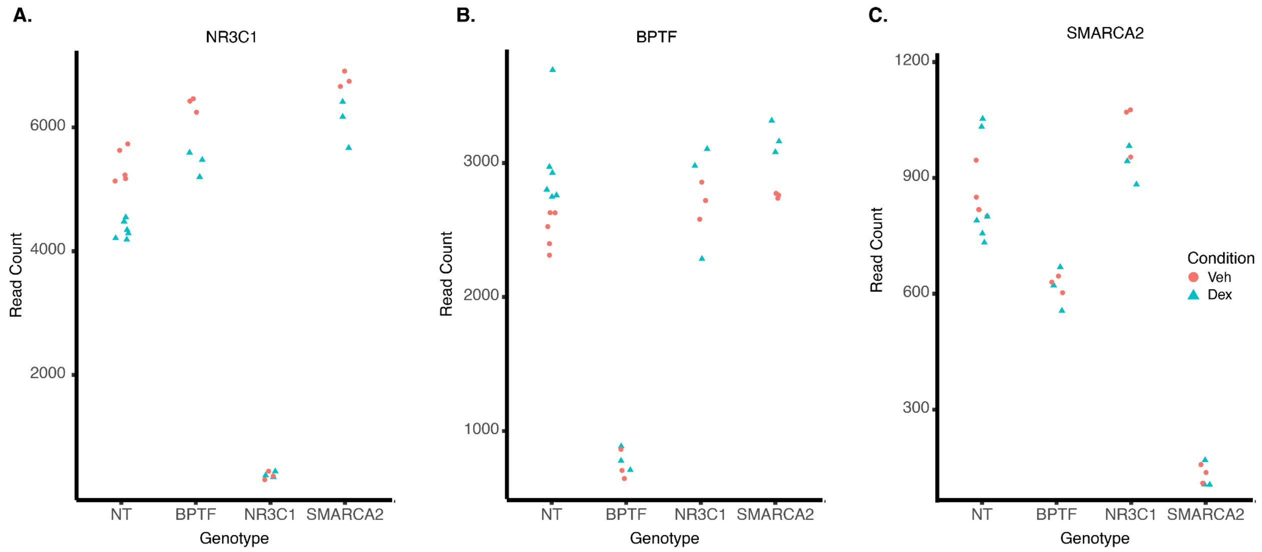


Normalized counts of gene transcripts across different siRNA-treated cell populations (n = 3 to n = 6 replicates per drug condition) for **A.** NR3C1, **B.** BPTF, and **C.** SMARCA2

### Supplementary Figure 3

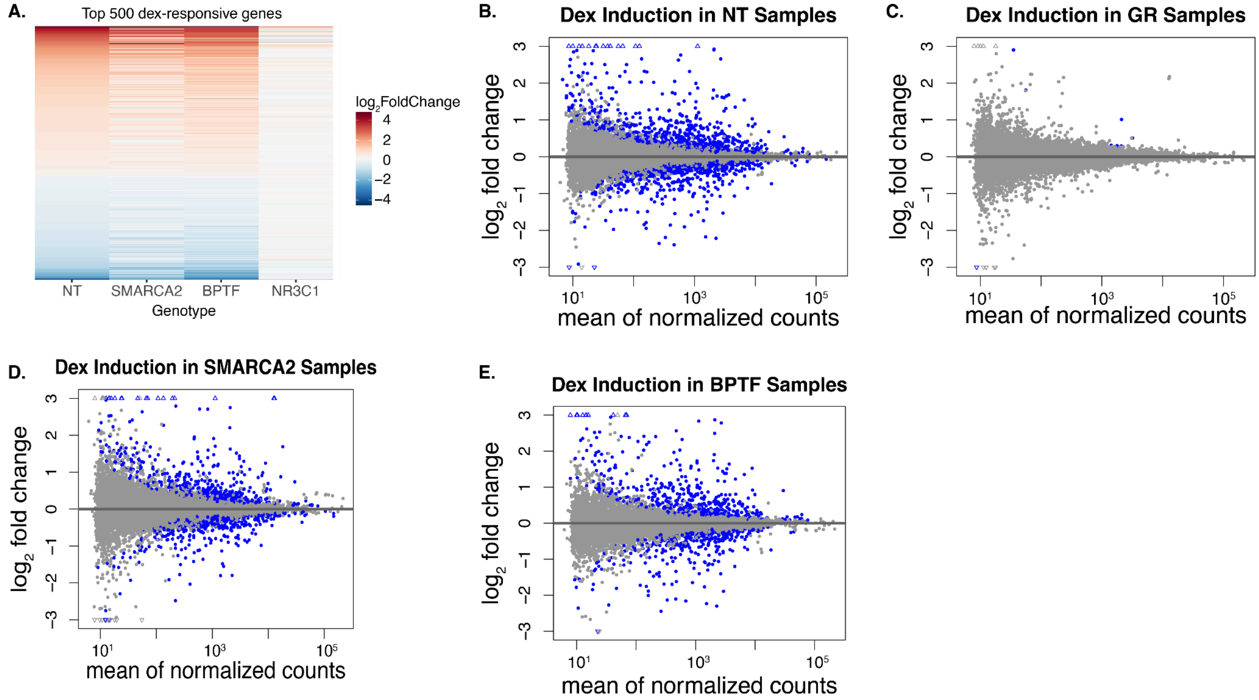


**A.** Heatmap showing the dex vs. vehicle $\log_2$ fold change change for the top 500 most significant dex-responsive genes for different siRNA-treated cell populations (n = 3 to n = 6 replicates for each siRNA) MA plots showing dex-induced gene expression changes in **B.** cells transfected with non-targeting siRNA, **C.** cells transfected with NR3C1-targeting siRNAs, **D.** cells transfected with SMARCA2-targeting siRNAs, and **E.** cells transfected with BPTF-targeting siRNAs

#### Supplementary Figure 4

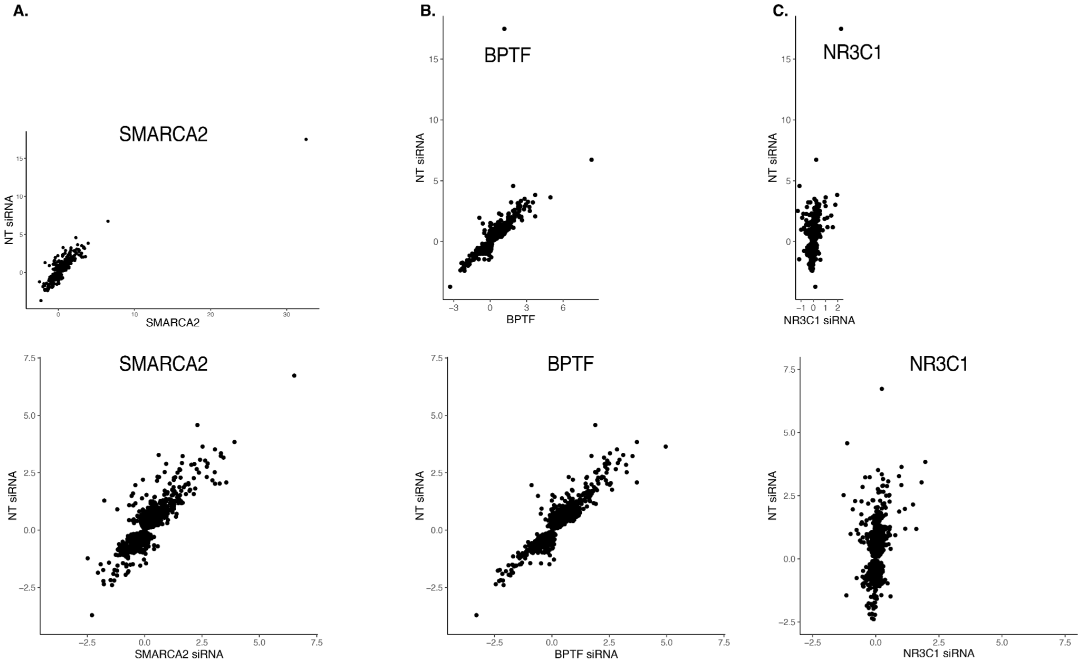


Correlations between the dex-induced $\log_2$ fold change in cells treated with the non-targeting siRNA and cells treated with siRNAs targeting **A.** SMARCA2 **B.** BPTF and **C.** NR3C1. The top panels show the full range of the data and the bottom panels show the data along the same scale with outliers removed

#### Supplementary Figure 5

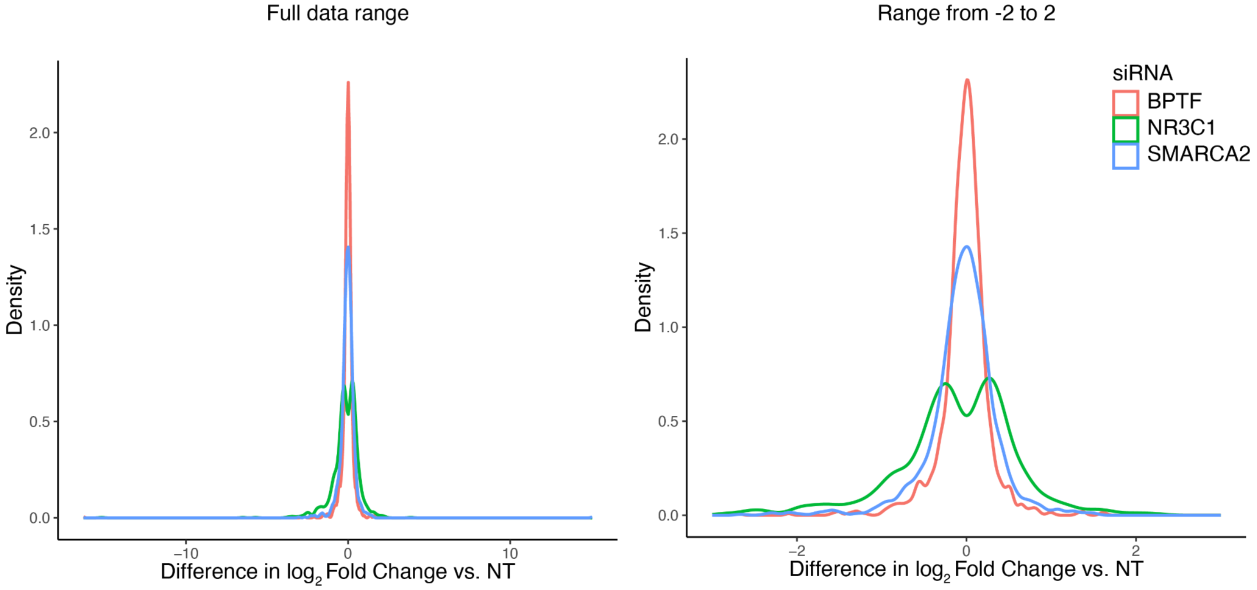


Density plot showing the difference in $\log_2$ fold change between cells treated with the non-targeting siRNA and cells treated with siRNAs targeting BPTF, NR3C1, or SMARCA2

#### Supplementary Figure 6

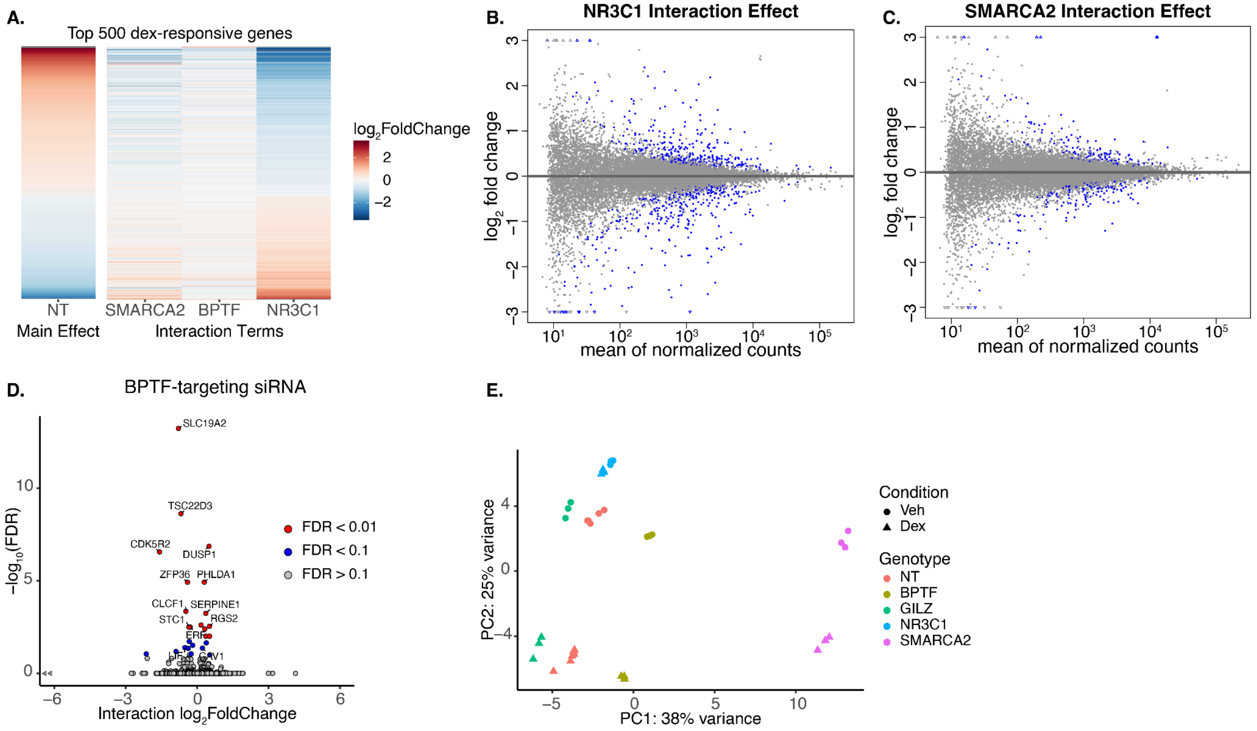


**A.** Heatmap showing the interaction $\log_2$ fold change for the top 500 most significant dex-responsive genes for different siRNA-treated cell populations (n = 3 to n = 6 replicates for each siRNA). Non-targeting siRNA column shows the main effect $\log_2$ fold change for reference. MA plots showing interaction $\log_2$ fold change changes for **B.** cells transfected with a NR3C1-targeting siRNA, **C.** cells transfected a SMARCA2-targeting siRNA, and **D.** cells transfected with a BPTF-targeting siRNA. **E.** Principal component analysis plot for the RNA Seq data across different siRNA and drug treatments.

### Supplementary Figure 7

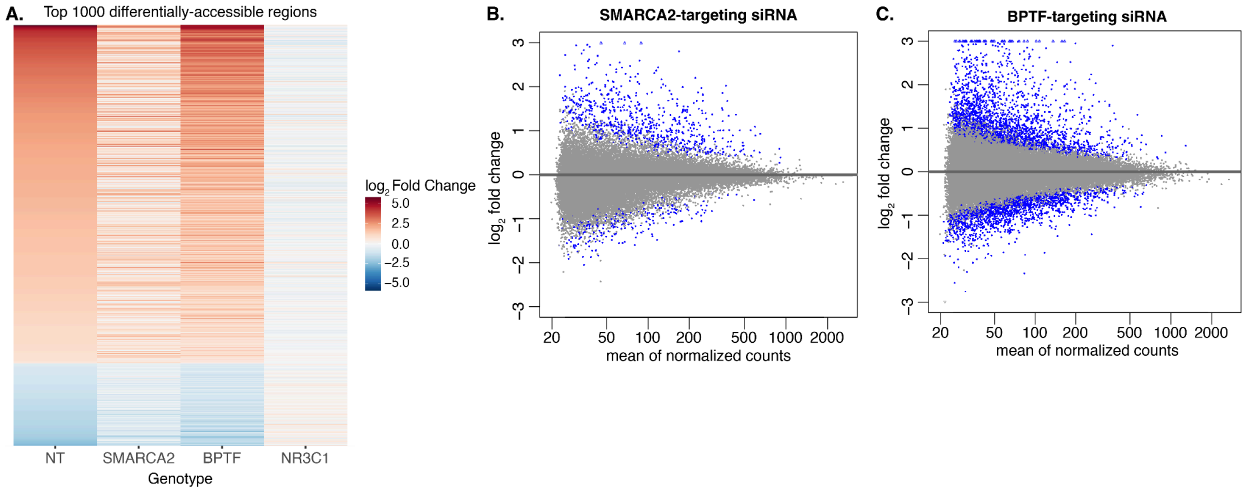


**A.** Heatmap showing the changes in chromatin accessibility as measured by ATAC-seq for dex treatment compared to vehicle for cells transfected with different siRNAs (n = 3 to n = 5 independent transfections per condition) **B,C.** MA plots showing dex-induced chromatin accessibility changes in **B.** SMARCA2 knockdown cells (n = 3 independent transfections) and **C.** BPTF knockdown cells (n = 3 independent transfections)

#### Supplementary Figure 8

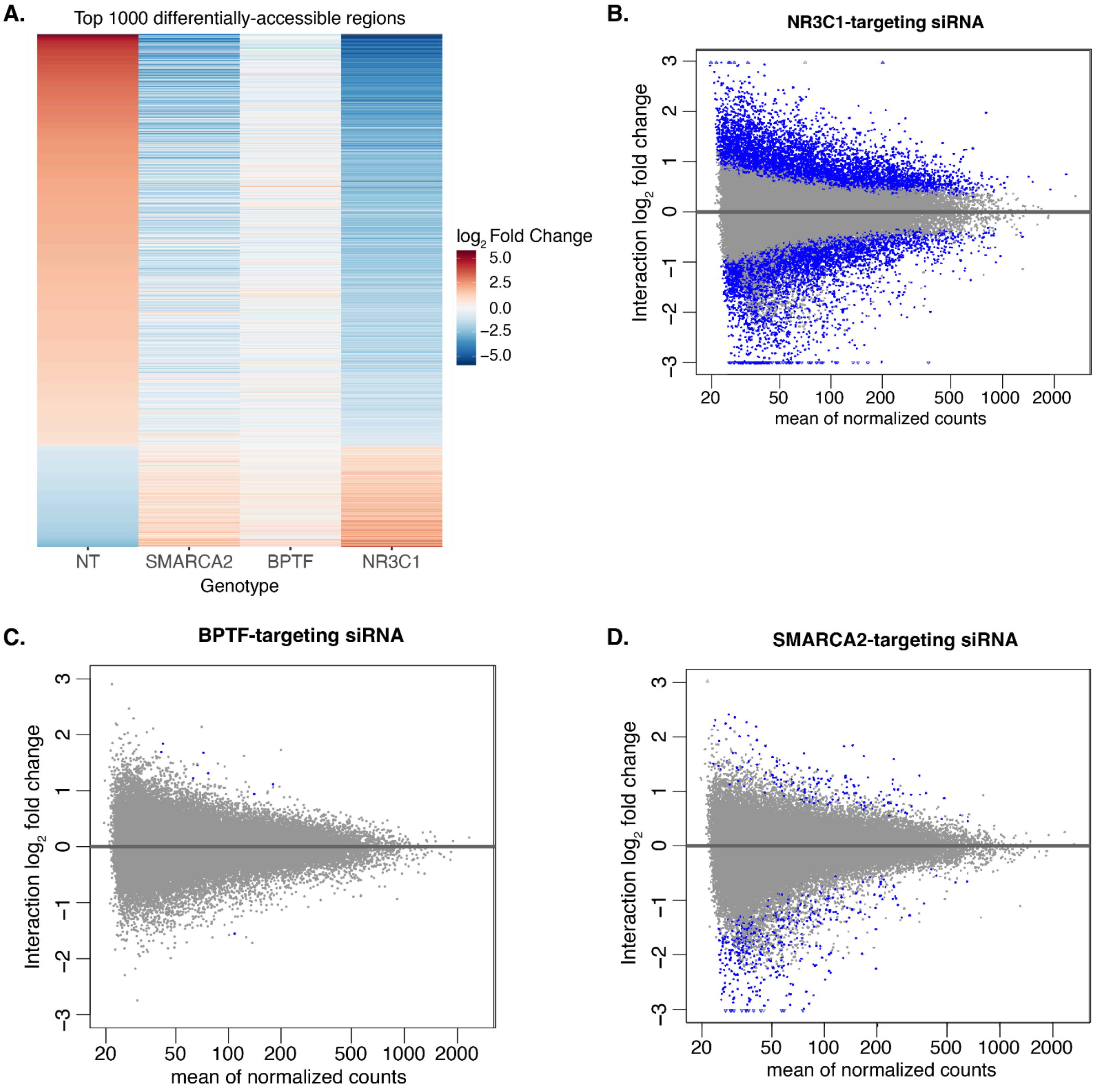


**A.** Heatmap showing interaction effects for dex-specific changes in chromatin accessibility as measured by ATAC-seq for cells transfected with different siRNAs (n = 3 to n = 5 independent transfections per condition). The NT column shows the dex-induced $\log_2$ fold change in the non-targeting condition for reference. **B-D.** MA plots showing interaction terms for dex-specific chromatin accessibility changes in **B.** NR3C1 knockdown cells (n = 3 independent transfections), **C.** BPTF knockdown cells (n = 3 independent transfections) and **D.** SMARCA2 knockdown cells (n = 3 independent transfections)

**Supplementary Figure 9**

**A.**

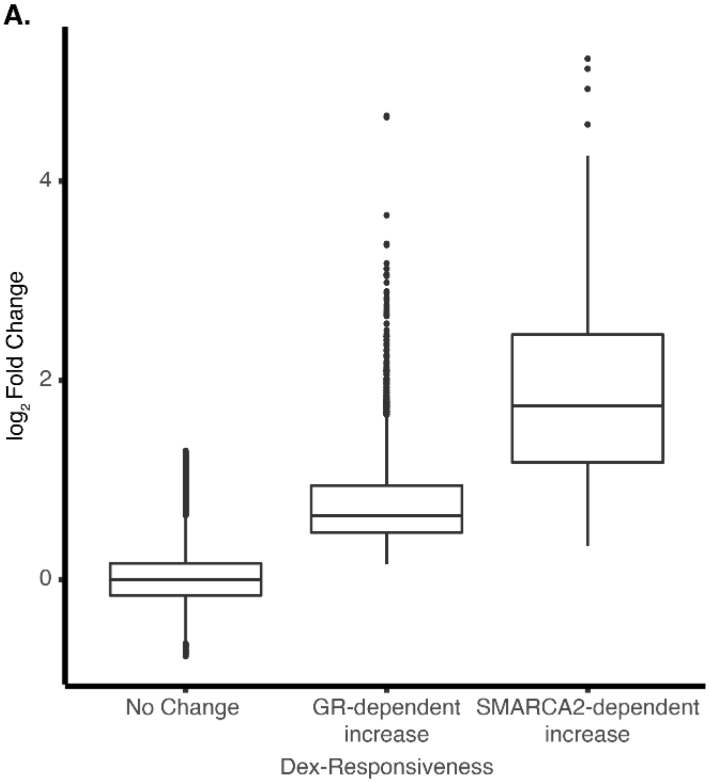


**A.** Boxplots showing the change in chromatin accessibility after dex treatment in A549 cells transfected with non-targeting control siRNA. Genomic regions are stratified by no change in accessibility, GR-dependent increase in accessibility, and SMARCA2-dependent increase in accessibility.

#### Supplementary Figure 10

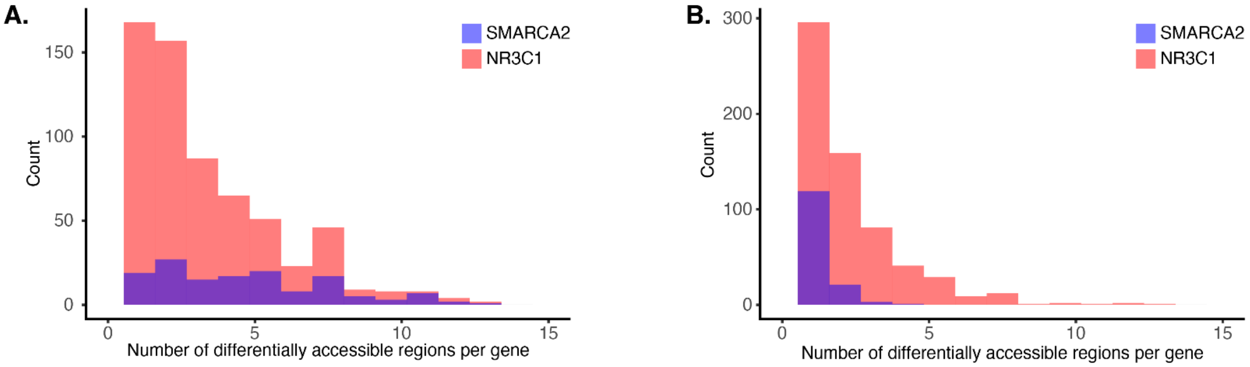


**A.** Histogram showing the number of genomic regions per dex-responsive gene with dex-induced differential accessibility. Blue denotes a gene associated with at least one genomic region with a SMARCA2-dependent accessibility change. Red denotes a gene associated with at least one genomic region with an GR-dependent accessibility change, but no genomic regions with SMARCA2-dependent accessibility change. **B.** Histogram showing the number of genomic regions per dex-responsive gene with SMARCA2 or GR-dependent differential accessibility. Blue denotes the number of regions per gene with a SMARCA2-dependent accessibility change. Red denotes the number of regions per gene with an GR-dependent accessibility change.

#### Supplementary Figure 11

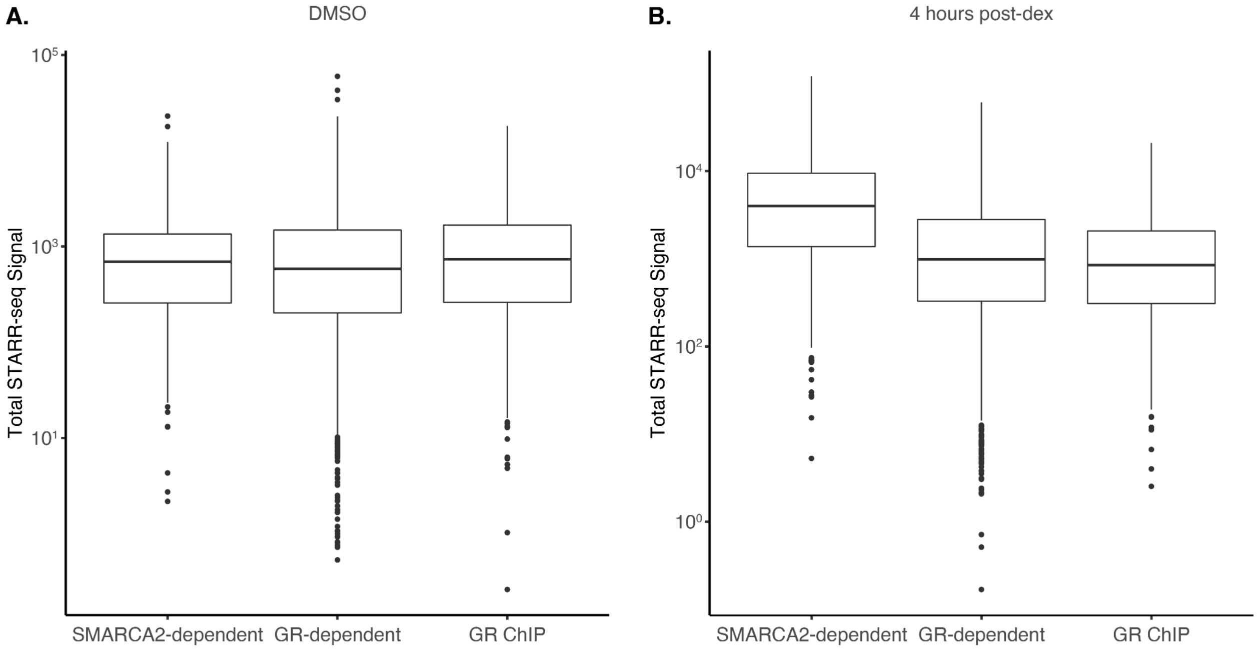


Box plots showing total STARR-seq signal at length normalized genomic regions with a SMARCA2-dependent accessibility change, GR-dependent accessibility change, or GR binding but no accessibility change after four hour treatment with **A.** DMSO or **B.** 1uM dexamethasone

#### Supplementary Figure 12

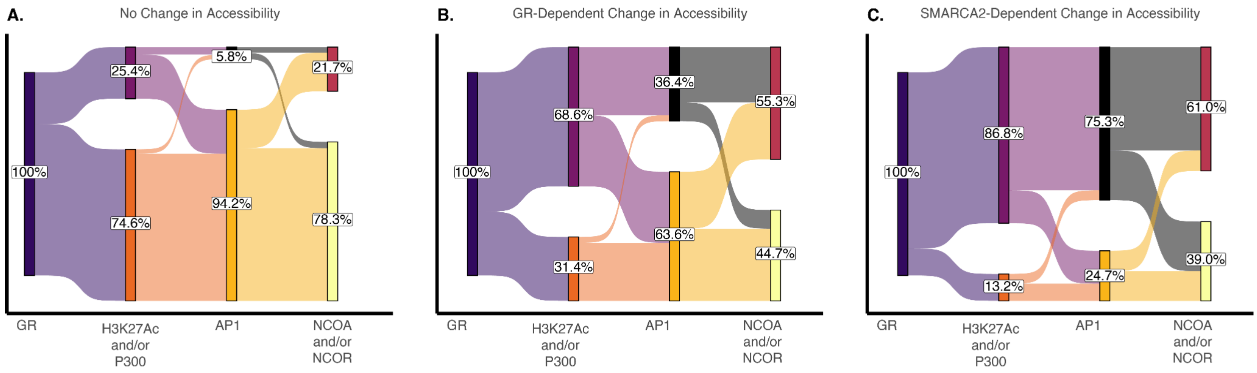


Sankey diagrams showing the proportion of genomic regions with positive differential transcription factor binding at 1, 4, or 8 hours after dex treatment for **A.** genomic regions with differential GR binding but no change in accessibility after dex treatment, **B.** genomic regions with differential GR binding and a GR-dependent change in accessibility after dex treatment, and **C.** genomic regions with differential GR binding and a SMARCA2-dependent change in accessibility after dex treatment

#### Supplementary Figure 13

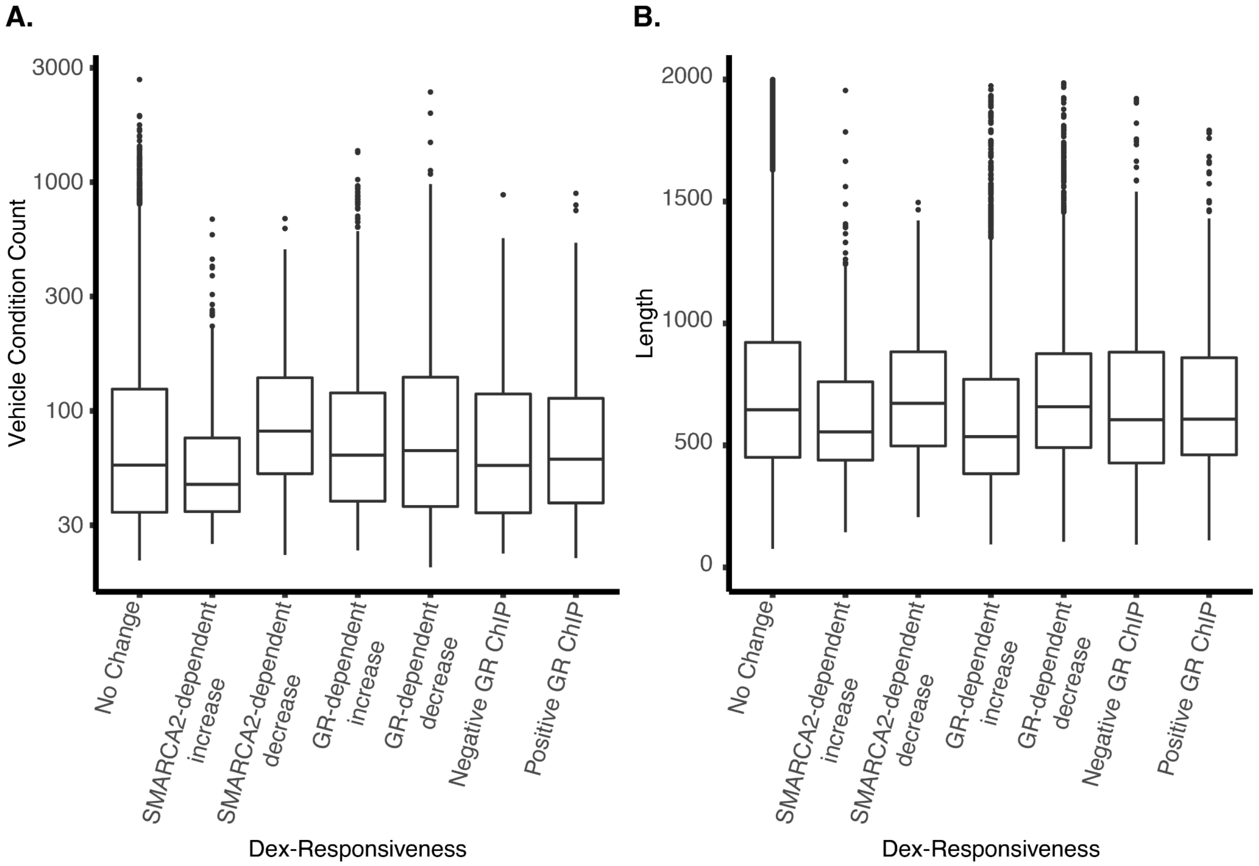


Boxplots showing **A.** The normalized number of reads in the vehicle condition in cells transfected with a non-targeting control siRNA or **B.** the length (in bp) for all regions of the genome identified via ATAC-Seq, stratified by whether they exhibit dex-induced changes in chromatin accessibility and/or differential GR binding after dex treatment
